# Genetic affinities and sub-structuring in Coorg population of Southern India

**DOI:** 10.1101/2022.07.20.500704

**Authors:** Anirban Mukhopadhyay, Lomous Kumar, Kiran Sran, Kumarasamy Thangaraj, B K Thelma

## Abstract

The Coorgs, also known as Kodavas, are one of the smallest religious and socio-culturally homogenous communities in the world, currently residing in the state of Karnataka, India. Due to a stark contrast with the surrounding subpopulations, their genetic architecture and population & demographic history have been a matter of debate for long. To better understand the population structure and demographic history of this caste group, we analysed the population, using high-resolution autosomal (n=70) as well uniparentally inherited markers (Y-chromosomal and mitochondrial DNA) (n=144). Our first ever findings elucidate that origin of Coorgs traces back to early or middle Bronze Age. We further found population substructure among Coorgs, which manifested as three distinct clusters in a Principal component analysis. One of these subgroups has undergone a rare and immense amount of population-specific drift but all three eventually admixed, both genetically and socio-culturally. The mtDNA analysis showed 40% South Asian-specific mitochondrial lineages among Coorgs; while the Y-chromosomal analysis revealed predominant presence of Eurasian, Middle-Eastern and Indian-specific haplogroups suggesting male-centric migration and eventual assimilation with local females. Our results for the first time identify these ancient and distinct genealogies that make up the contemporary Coorgs and may explain the socio-cultural differences with their immediate and distant neighbours in the country and the prevalent narrative history. In a wider context, the study also reveals an ancient, yet unknown, Middle Eastern source population that might have contributed to an early west to east migration into India.

## Introduction

The population history of India is rich, varied, interesting as well as intriguing. The extensive diversity in this rather old subcontinent is due to multiple waves of migration into the region from foreign lands, over millennia and eventual geographical and linguistic isolation and/or prevalence of caste endogamy. The country continues to be popular for explorations of genetic architecture of ancient/isolated human populations with the most recent being the novel and exciting findings from the Rakhigarhi excavation sites (Shinde, et al. 2019). Furthermore, India with its diverse geological landscapes ranging from plateaus, hills to dense forests offered sub-populations a chance to occupy niches, leading to further isolation. The Deccan plateau, carpeting a large part of Southern India, came into being in the Cretaceous period and now extends over eight states of modern India (Eaton, et al. 2005). Historically, it witnessed the rise and fall of some of the most momentous and longest standing dynasties of not only India but also of the world. It was the cradle of ancient Mauryan empire (300 BCE), the earliest recorded, and home to powerful medieval dynasties such as the Cholas (3rd century BCE to 12th century CE), the Chalukyas (6^th^-12^th^ century CE) and the subsequent, modern Vijayanagara empire (1336–1646) (Rao 2005). Of note, the Deccan saw its own share of vassals from distant lands and immigrants (Thapar 2015), similar to the peopling of north India (Joseph 2018). Thus, since antiquity the plateau has been a hotbed of civilization, culture and learning. The resultant of such socio-cultural variations together with caste endogamy and/or geographic isolation led to the emergence of some of the contemporary finer populations.

One such group, the Coorgs (also referred to as Kodavas) inhabit the district of Kodagu (Coorg) in the Southern Indian state of Karnataka, nestled away in the Western Ghats, the geographically isolated, eroded slopes of the Deccan Plateau. The term Kodava serves a dual purpose encompassing the language and culture and also denotes the dominant community of the region, who inhabited the land from pre-historic times. Traditionally a group of agriculturists with martial customs interlacing their day-to-day lives, the Coorgs have continued to practice family exogamy and caste endogamy till date (Cariappa and Cariappa 1981). Furthermore, the Coorgs are distinctly different from the neighbouring populations both in terms of religious as well as socio-cultural practices. Though they follow Hinduism, various customs prevalent among the Coorgs notably deviate from the Hindu way of life (Bopanna 2022). These include i) Caste system, unlike in Hindus is nearly non-existent amongst the Coorgs; ii) the Brahmin, so called supreme caste, is not revered in their rituals and ceremonies related to birth, coming of age, marriage, and death but are presided over by the elders of their community; iii) Ancestor and nature worship is prevalent with each Coorg belonging to a patrilineal okka or clan and are the descendants of a common ancestor known as the Karanava, worshipped akin to a Godhead; and iv) The religious dynamics involve an ancestral home (Ainmane) with a small shrine (Kaimada) dedicated to the ancestors of the respective clans; a lamp (bolcha) or a hanging lamp (thookbolcha) lit in the central hall (nellakki nadubade) of the ainmane at dawn and dusk. The offerings (meedi) made in honour of the ancestors usually include besides other food items, pork and liquor which are largely forbidden among Hindus. These deviations have led to the two major models of the origin of the Coorgs.

### The native model

The indigenous model holds many views but primarily ascribes the Coorgs as pre-historic inhabitants of the Kodagu region. One view based on the measurements of cranial and nasal index is that the Coorgs are the descendants of the brachycephalic humans, who entered the Indus Valley during the Mohenjodaro period (before the immigration of the Indo-Aryans) and sometime later migrated to the Kodagu region (Hutton as quoted by Balakrishnan 1976) (Balakrishnan 1976). The Kaveri Purana an inset of the ‘Skanda Purana’ (8^th^ Century CE) classifies the Coorgs as a warlike native tribe of Kodagu, who learnt and started practicing agriculture. It also mentions Chandra Varma, a king of the ancient lunar dynasty, who came across the region on one of his expeditions, became the king of the land and settled there with the natives.

### The non-native model

This model ascribes Coorgs as the descendants of tribes or clans that immigrated into India at different time periods. These theories often levy Coorg lineage to i) the Indo-Greeks who remained in India after the conquests of Alexander; ii) the pre-muslim Kurds or pre-Christian Georgians escaping religious radicalization (Ponnappa 1999); and iii) being an off-shoot of the Indo-Scythian Sakas (Connor 1870; Rice 1877). These opinions lack scientific evidence and are mostly based on the various parallels found in the socio-cultural dynamics including attire and folk practices of the Coorgs with that of the respective foreign populations. With most views for the origin of the Coorgs thus being entirely anecdotal or from the early years of population dating, evidence-based understanding as to their origin is notably lacking. However, historians agree that Coorgs have been the earliest inhabitants of Kodagu for over thousands of years (Bowring 1872; Kamath 1993; Rajyashree 2001).

Genetic markers and genome analyses tools have been instrumental to uncover intriguing population histories of immense scientific and public interest over the last three decades. In this study, using high-resolution mitochondrial DNA (mtDNA), Y-chromosomal and autosomal genetic markers on a representative Coorg sample set, comprised of only males i) the incumbent genetic architecture; ii) the likely time of their origin; and iii) inter-population relatedness across contemporary and ancient global populations were investigated. This first report, has demarcated the socio-culturally homogenous present day Coorg population into three distinct sub-groups (Coorg1, Coorg2 and Coorg3) with two of them being ancient populations. Coorg1 belongs to the Indian subcontinent with high ASI ancestry and Coorg3 is with high early-Bronze Age Namazga ancestral component but both with considerable genetic drift. Coorg2 on the other hand, though similar to its sub-continental neighbours (ANI), showed signs of admixture with Coorg1 and Coorg3. These results highlight a genetic as well as socio-cultural admixture of the three distinct and early groups lending to the formation of the present day Coorgs.

## Results

### Autosomal markers

#### Substructuring in Coorg population

Principal component analysis (PCA) was performed to gain insight into population structure. Interestingly, the Coorg comprised of three distinct genetic clusters, we named as Coorg1, Coorg2 and Coorg3, which, were placed far away from each other (Fig 1A). Coorg1 was clustered near one extreme of the south Asian cline where most of the diverged Dravidian groups with highest ASI ancestry were found. Coorg2 showed affinity with Indo-European population groups and to additional groups like Nair, Bunt, Hoysala, Reddy, Thiyya and Ezhava, with higher Middle Eastern component (Kumar, et al. 2022). Coorg3 individuals were not clustering with any of the modern Eurasian or south Asian populations. Rather this unique cluster was placed near the extreme top left of the plot, unlike South Asian and all other modern Eurasians, who were at the bottom of the plot as a single continuous cline of high West Eurasian to low West Eurasian affinity (Fig. 1A). Second group did not show complete affinity to Indo-Europeans in South Asian cline but was slightly shifted in the direction of the third group characterised by a completely isolated cluster, thus suggesting their possible affinity with Coorg3.

**Fig. 1.**
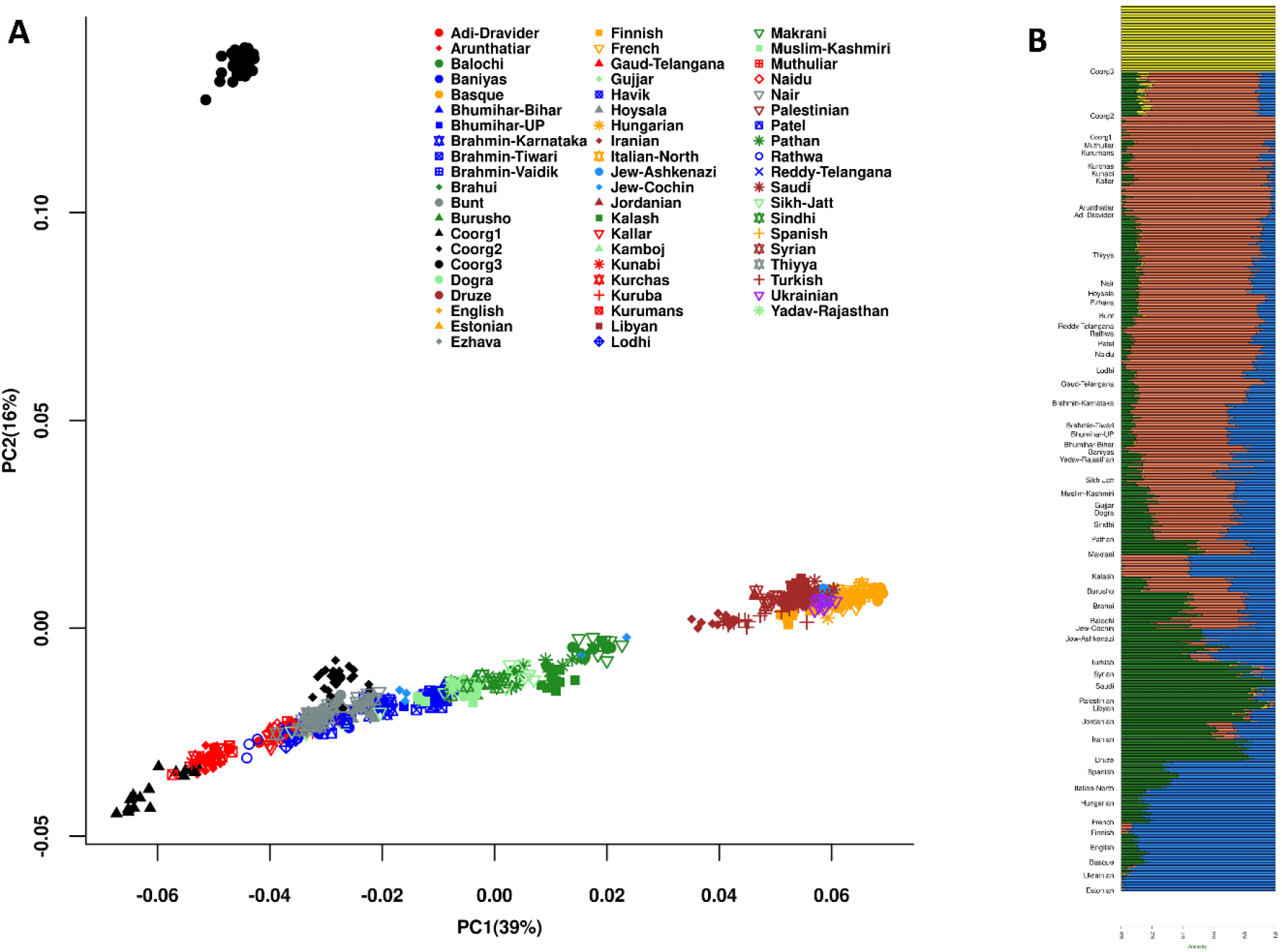
PCA and ADMIXTURE Analysis. **A**. Biplot of principal component analysis (PCA) of Coorgs with modern Eurasian populations with first two components. B. Stacked barplot of the ADMIXTURE analysis with K=4 with global populations ordered geographically.

#### Unique ancestral component of Coorg3 in ADMIXTURE analysis

In order to infer the ancestral genetic components in the context of modern Eurasians and to further inquire about the clustering pattern found in PC analysis, we used model-based approach in ADMIXTURE with K=4 (Fig S1). Surprisingly, Coorg3 population, which was completely isolated in PCA clustering, formed a unique yellow component different from other Eurasian population groups (Fig. 1B). Coorg2 was similar in ancestry profile to Indo-Europeans (ANI), as they had similar proportions of blue, red and green components with the only difference of having an additional yellow component, a feature of Coorg3. Coorg1 individuals were enriched for the South Asian-specific red component.

#### Admixture history confirms affinity of Coorg1 with ASI and Coorg2 with ANI

Admixture history and allele sharing pattern were further tested in the three groups by utilizing admixture F3 and Dstatistics methods in qp3Pop and qpDstat tools of ADMIXTOOLS (Patterson, et al. 2012) package. Admixture F3 was run in the form F3 (X, Palliyar; Coorg1/Coorg2/Coorg3) using Palliyar as proxy for ASI source of ancestry and population X as different West Eurasians and South Asian populations. Coorg3 did not have any signal of admixture, with a complete absence of negative admixture F3 statistics with any combination of source groups (supplementary table_1a). Coorg1 was also devoid of any such signal of admixture (supplementary table_1b). Consistent signal of admixture was found only with Coorg2 individuals, which showed significant admixture with all West Eurasian groups tested and in South Asia with groups having higher ANI component like Kalash and Pathan from Pakistan and Sikh_Jatt, Kamboj, Dogra, Bhumihar_Bihar, Brahmin_Haryana, UP_Brahmin, etc. from North west and North India (supplementary table_1b). They also had significant Admixture F3 statistics (Z score = -7.037) with Coorg3 individuals, indicating some shared ancestry in the past (supplementary table_1c). Dstatistics in the form of F4 (X, Coorg1/Coorg2/Coorg3, French;Yoruba) and F4 (X, Coorg1/Coorg2/Coorg3, Palliyar; Yoruba) were compared to infer the placement of the three groups with respect to relative ANI and ASI affinity in context of other Eurasian groups (supplementary table_1d-f). In the scatterplot, Coorg1 showed highest ASI affinity outcompeting all other south Asian or west Eurasian groups (Fig S2A). It also showed negative D-statistics in the run in the form F4 (X, Coorg1/Coorg2/Coorg3, Palliyar; Yoruba) with all the West Eurasian or South Asian groups, giving some indication of sharing same clade with Palliyar (supplementary table_1d). Coorg2 was outcompeting some of the Indo-Europeans with lesser ANI component and all Dravidian and Austroasiatic groups in terms of ANI affinity, an indication of moderate ANI to ASI ratio (Fig S2B). Coorg2 and Coorg3 showed similar affinity patterns, but Coorg2 slightly outcompeted Coorg3 in ASI and ANI affinity (Fig S2B). But this lesser affinity may be the consequence of more drift in Coorg3 as was observed in F3 admixture test (supplementary table_1c).

#### Genetic dissection of the three groups uncovers ancient ancestral lineages

Distal and proximal modelling approach was applied with early-Bronze age and Bronze age sources respectively, using qpAdm of ADMIXTOOLS to compare the ancient ancestral contributions in the three groups of Coorgs and a few other South Asian populations. In distal modelling, Andamanese Hunter-Gatherers (AHG), Chalcolithic Namazga samples (Namazga_CA) and Middle or Late Bronze age Steppe groups (Steppe_MLBA) were used as left groups. Among all South Asian groups tested, Namazga_CA component was comparatively higher in proportion in Coorg3 (0.46), second after Gujjar (0.53) and similar to Kamboj (0.46) followed by Pathan (0.45), Nair (0.44) and Coorg2 (0.44) (Fig 3a). Interestingly, Coorg1 had similar composition of AHG (0.7), Namazga_CA (0.2) and Steppe_MLBA (0.05-0.06) to Palliyar (Fig 2A) (supplementary table_1g).

**Fig 2.**
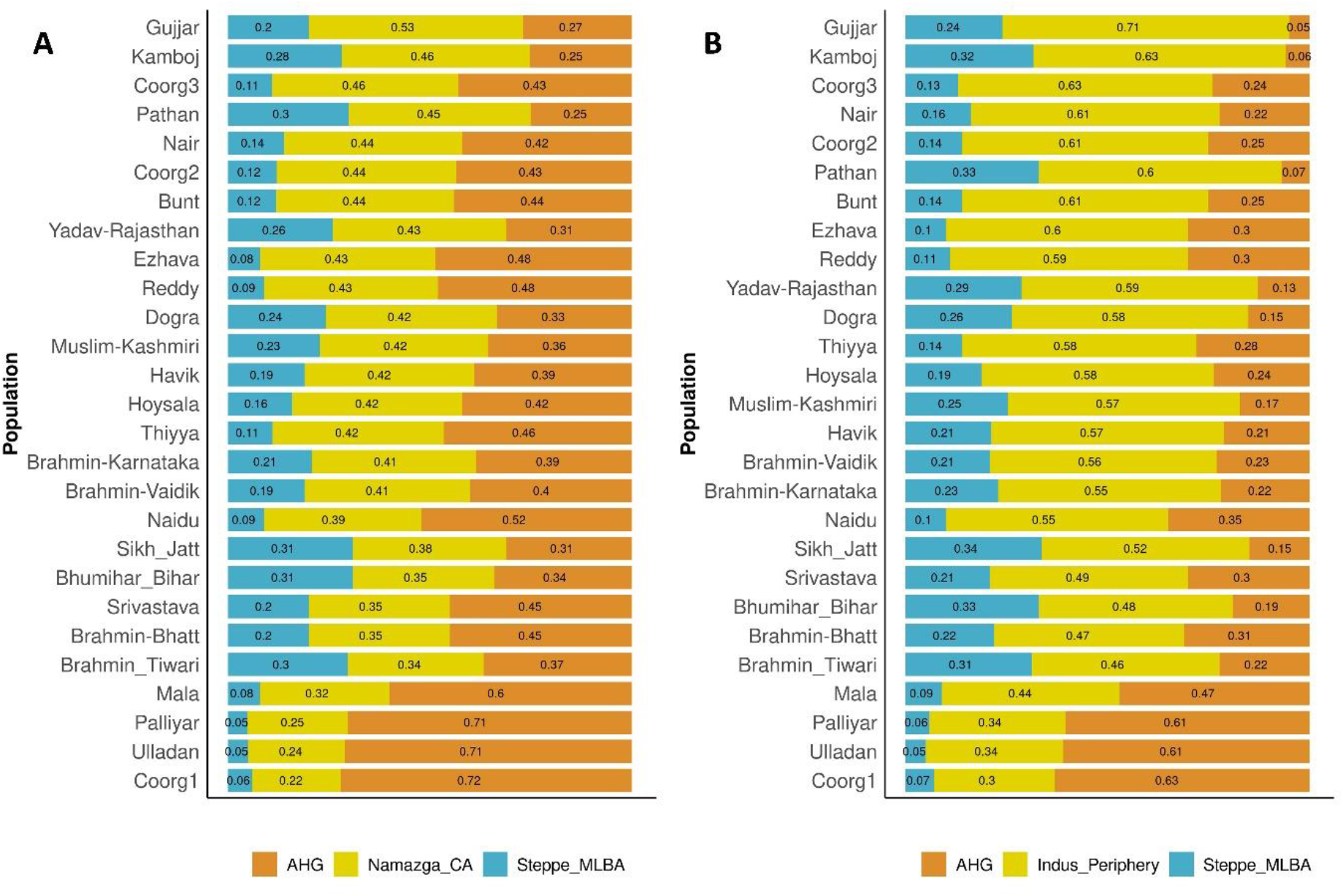
A. Distal and B. Proximal admixture modelling with qpAdm for three Coorg groups and other Indo-European and Dravidian populations.

**Fig 3.**
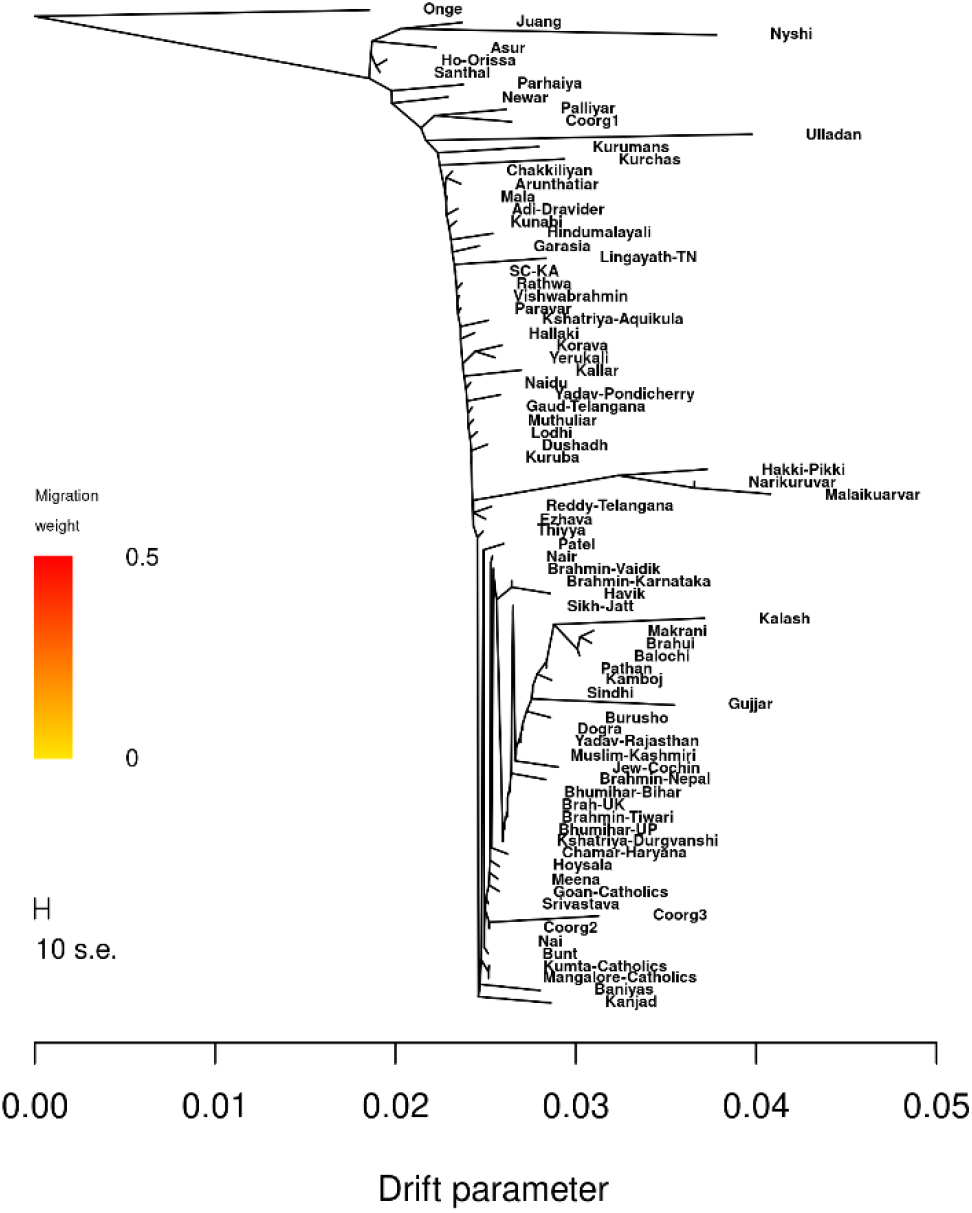
Maximum likelihood tree constructed using TreeMix to infer association with Indian Indo-European caste groups and population-specific drift.

In the proximal modelling with Bronze Age sources (AHG, Indus_Periphery and Steppe MLBA), Coorg3 had second highest contribution from Indus_Periphery group jointly with Kamboj (0.63), next to Gujjar (0.71), followed by Nair (0.61) and Coorg2 (0.61) (Fig 2B) (supplementary table_1g).

#### Clues towards an ancestry from a ghost lineage for Coorg3 from qpGraph

The three Coorg groups were jointly fitted into the Admixture graph topology using qpGraph implementation of Admixtools 2. Initial graph based on previous South Asian graph (Narasimhan, et al. 2019) was used as a starting point with some simplification and automated graph exploration was run 30 times for each group using *find_graph* feature of Admixtools 2 (Maier, et al. 2022). Of the 10 best fits with score closer to zero and consistent with the known admixture history of South Asia, the top two best fitted topology are shown. First graph topology is with likelihood score 2.032829 (worst Fstat z-score 0.9738), while second graph is having likelihood score of 2.417081 (worst Fstat z-score 1.0144). Both graph topologies were compared in terms of out of sample score and bootstrap resample scores and it was found that both graphs performed well and similar, with the second graph having only a slight advantage over first (supplementary table_1h). But it was observed that both graph topologies required an additional source of ancestry from a ghost lineage for Coorg3 group with 1% and 9% contribution for first and second graph, respectively (Fig S3A-B). Same was true with all the 10 best fitted graph topologies.

#### Extreme population-specific drift and divergence in Coorg3 subgroup

In the Maximum likelihood tree constructed using TreeMix v.1.12 (Pickrell and Pritchard 2012), it was found that although placed among Indian Indo-European caste groups, Coorg3 shows significant amount of population-specific drift as indicated by longer branch length compared to other similar groups in South Asia (Fig 3). In South Asia only Kalash and Gujjar had more population-specific drift than Coorg3. Coorg2 was also placed in same clade with Coorg3, however, not with significant drift. Interestingly, Coorg1 shared the same clade with Palliyar with exactly similar amount of drift and both shared the clade with Ulladan, which had a much higher amount of drift (Fig 3).

In the Wier-Cockerham F_st_ measurement across the genome for three groups of Coorg population, surprisingly higher distribution of F_st_ value for Coorg3 was found, with chromosome 19 and chromosome 22 having the greatest number of SNPs with highest F_st_ values. This was true with all three comparison groups viz, Dravidian, European and Middle East (Fig S4G-I). For Coorg1 and Coorg2, the FST values for SNPs were in normal range as observed with other groups. But Coorg1 and Coorg3 showed similar amount of drift as that of Austroasiatic group in distance matrix of WC-Fst (Fig S5). The extent of population-specific drift among Coorg2, Coorg3, Kalash and Gujjar were compared using qpGraph. Coorg3 had similar amount of drift as that of Kalash and Gujjar, while Coorg2 did not have any significant population-specific drift (Fig S6A).

#### Fine scale population structure and haplotype sharing

To gain a better understanding of population structure and haplotype sharing pattern of the three Coorg groups with modern Eurasians, haplotype-based approach with ChromoPainter (Lawson, et al. 2012) and fineSTRUCTURE (Lawson, et al. 2012) was used. The PC analysis with the co-ancestry matrix computed by ChromoPainter clearly differentiated Coorg3 from other populations with distinct clustering at top right corner, while all other groups were forming a cline along the diagonal (Fig S7). Some individuals of Coorg2 were in the main Indian cline, while most of them were inclined towards Coorg3. There was a split in the Coorg1 cluster too, with one group placed along with Ulladan and Palliyar, while another group completely separated and was placed at the extreme end of the Indian cline. One of the individuals from Coorg2 moved in the cluster of Coorg3 (Fig S7).

fineSTRUCTURE kept Coorg3 individuals in a separate cluster of 9 clades with other West Eurasian and South Asian groups, which were in two separate clusters of 6 and 29 clades, respectively (Fig S8-9). South Asian cluster was further subdivided into two sub branches. One of the branches was of main cluster of Indo-European and Dravidian caste and tribe groups, while other cluster was of relatively diverged groups of Onge, Austroasiatic, Tibeto-Burman and Dravidians. First main cluster had Coorg2 individuals placed between Indo-Europeans and Dravidians. Coorg1 individuals were placed among second South Asian cluster of diverged groups, which was further subdivided into three clusters. One of these was with Onge, Nyshi and Chakehshanega, second cluster was with most of the Coorg1 individuals and a single Kuruman individual and third cluster was with the remaining Coorg1 individuals along with Ulladan, Palliyar, Kurchas and Kurumans (Fig S8-9).

#### Runs of Homozygosity, relative IBD score and Admixture dating

In the Runs of Homozygosity (RoH) analysis using Plink 1.9 (Chang, et al. 2015) Coorg1 showed highest mean of total length of RoH at the window size of 1000kb, outcompeting even Palliyar, Kurchas, Kurumans and Kalash (Fig S10A). But in terms of mean total number of RoH, Palliyar had the highest value for distribution. Interestingly, Coorg3 had the least distribution in terms of both mean length and mean number of RoH among all the South Asian populations. Coorg2 was placed along with other Indo-European caste groups like Brahmin_Tiwari, Dogra and Lodhi indicating higher effective population size and low level of recent consanguinity. But for higher window sizes of 2500kb (Fig S10B) and 5000kb (Fig S10C), Palliyar, Kalash and few other groups like Baluchi, Kurchas and Kallar were taking over Coorg1 in terms of overall distribution of RoH.

IBD score relative to Finnish population for Coorg1 was very high (2.854) compared to Coorg2 (1.447) and Coorg3 (0.6), and was very close to Vysya (3.122), Reddy_Telangana (2.19) and Panta Kapu (2.175) (Fig S11) (supplementary table_1i).

The approximate date of admixture of west Eurasian and South Asian genetic components in the genomes of the three groups was tested using linkage disequilibrium-based method implemented in ALDER. Juang population was used as proxy for South Asian ancestry and a different west Eurasian population as another reference group. Coorg3 showed admixture signal with all west Eurasian groups (2-ref Z score > 5), which dates approximately to ∼70-80 generations before present (supplementary table_1j) (Fig S12D-F). But for Coorg2 distribution of West Eurasian admixture date was comparatively earlier (95-100 generations before present) with more significant P_val and 2-ref Z score (supplementary table_2k) (Fig S12A-C). Run with Coorg1 was only successful while using Palliyar as one source but those were not significant (2-ref Z score < 5) (supplementary table_2l).

#### Population separation history and Y STR-based network analysis

Genome-wide genealogy approach implemented in Relate was used to infer population separation history of the three groups. It was found that the separation of Coorg3 from other modern populations like Europeans, Middle-East, North West India, North India and Dravidian occurred much earlier than any of these groups from each other (i.e., middle or late Bronze age separation) (Fig S13C). Time of population separation of Coorg2 coincides with other Indo-European and Dravidian groups from India (Fig S13B), while Coorg1 showed an earlier separation with a population bottleneck after separation (Fig S13A).

Median-Joining (MJ) network for two sister clades R1a-M780 and R1a-Z2125 was constructed using a dataset of 17 Y chromosome STRs of the three Coorg groups. Samples not used for genome-wide SNP array (for whom affiliation to Coorg1, Coorg2 and Coorg3 is unknown) were also included. In the analysis with both markers (M780 and Z2125), distinct clustering of Coorg individuals in a separate branch and not with any of the Central Asian, Middle East or South Asian branches was observed (with exception of a single individual with Central Asians in case of M780) (Fig 4A-B).

**Fig 4.**
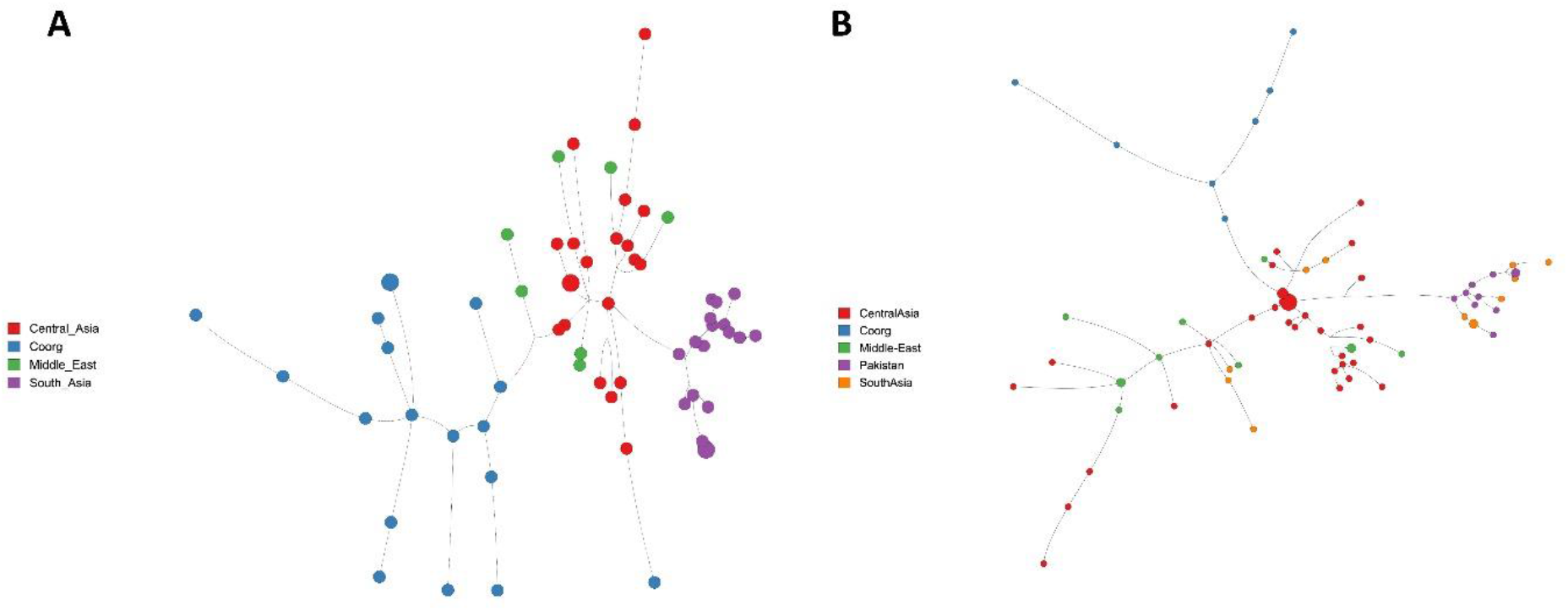
Median-joining Y-STR network for M780 and Z2125

#### Mitochondrial markers enriched for South Asian haplogroups across the three Coorg groups

Mitochondrial haplogroup diversity was further compared among the three subgroups. Coorg2 and Coorg3 were more diverse compared to Coorg1 in terms of haplogroup diversity, with presence of four major haplogroups (M, U, R and H or HV) (supplementary table_2a-c). All the three Coorg subgroups had highest frequency of South Asian-specific major haplogroup M, with a frequency of 0.54 in Coorg1, 0.38 in Coorg2 and 0.48 in Coorg3. The second most abundant mtDNA haplogroup was H, with a frequency of 0.23 in both Coorg1 and Coorg3 and 0.25 in Coorg2. Haplogroup U was observed only in Coorg2 (0.25) and Coorg3 (0.16) subgroups, while mtDNA haplogroup R was present with highest frequency in Coorg1 (0.15), followed by Coorg2 (0.08) and Coorg3 (0.07). Apart from these major haplogroups, unique haplogroups such as haplogroup G in Coorg2 (0.04), haplogroup HV in Coorg3 (0.07) and haplogroup T in Coorg1 (0.08) were also observed (supplementary table_2a-c).

In comparison to mtDNA haplogroups, more drastic differences in terms of Y chromosomal haplogroup distribution were observed (supplementary table_2a-c). There were no large differences in terms of distribution of middle eastern J haplogroup and Indus civilisation related L haplogroup, with both present with comparable frequency in Coorg2 (J=0.18, L=0.14), Coorg3 (J=0.19, L=0.16) and Coorg1 (J=0.29, L=0.14). However, South Asian-specific Y chromosome haplogroup H was observed with highest frequency in Coorg 3 (0.45), lesser in Coorg 2 (0.14) and with a complete absence in Coorg 1 (see discussion). Steppe related haplogroup R1 was highly prevalent in Coorg1 (0.5), followed by Coorg2 (0.32) and Coorg3 (0.07). South Asian-specific haplogroup R2 was observed in all three groups with moderate frequency. Some unique haplogroups such as haplogroups O in Coorg2 (0.04) and Coorg3 (0.03) and haplogroup K in Coorg2 (0.04) were also observed.

## Discussion

The Indian population is characterised by a number of high to moderately isolated populations, which pose a stark contrast to their immediate as well as sub-continental neighbours due to a plethora of geographical, cultural and linguistic factors. With only anecdotal accounts of their ancestry and origins, these populations offer a unique opportunity to anthropologists and geneticist to understand their genetic history with consequent unravelling of erstwhile unknown human migrations a welcome happenstance. This study is on one such isolated and small population of Coorgs, socio-culturally unique, early inhabitants of Kodagu district in Karnataka. Current knowledge of their migration history is also largely anecdotal or limited to cultural comparisons across global populations and population history is highly debated. This first in-depth genetic analysis of the Coorgs revealed their ancient origin, immense population drift due to isolation and a notable contrast with its neighbouring populations.

Analyses using mitochondrial, Y chromosomal and genome-wide autosomal markers along with contemporary statistical tools revealed an interesting delineation of this otherwise religious and socio-culturally homogeneous present day Coorg population into three distinct groups that we named as Coorg1 (C1), Coorg2 (C2) and Coorg3 (C3) (Fig1). Not unexpectedly, ADMIXTURE analysis revealed a moderately shared genetic component between C1 and C2; and C2 and C3 but negligible between C1 and C3 (Fig 2). The significant negative F3 statistics amongst these groups further re-iterates this observation. To cement this genetic admixture, in the co-ancestry matrix generated using Chromopainter and fineSTRUCTURE (Fig S8), C2 cluster was heading towards C3 indicating some affinity to C3 (Fig S9). The distinct clustering of C1 and C3 is supported further by the values observed in runs of homozygosity approach (Fig S10a-c). It is evident from the results that C1-C3 divergence pattern is not because of recent consanguinity or high-level inbreeding. C1 had higher level of RoH measures, almost comparable to those of old population groups like Palliyar, Kallar and other divergent groups (Fig S19A-C). Higher relative IBD score of Coorg1 further indicates the presence of endogamy and founder event in Coorg1 compared to other two groups (Fig S11) (supplementary table_1i). On one hand higher mtDNA diversity observed among C2 and C3 reflects the past expanding nature of these populations whereas C1, with its more or less homogenous haplogroup distribution indicates a population bottleneck, as also reflected in the autosomal analysis. High frequency of Y chromosome haplogroup R1 in C1 may also be a consequence of an identical founder event in the C1 population. This suggests that C1 may have suffered similar kind of drift as that of Palliyar due to long term isolation (discussed later).

At this point, it is important to discuss the genetic architecture of C2 and C3 in the light of their neighbouring as well as global ancient and contemporary populations. Mitochondrial DNA diversity of all the three Coorg groups reflects their mostly South Asian-specific maternal lineage along with some West Eurasian admixture owing to the presence of haplogroups HV, H, U and T with noticeable frequency, However, in the frequency based PCA analysis (Fig 1), C2 showed moderate affinity to the sub-populations with ANI ancestry such as Nair, Bunt, Thiyya and Hoysala, which have been reported to have comparably higher Middle Eastern components. Comparatively higher frequency of haplogroup R1 in Coorg2 reflects their more local admixture with Indo-Europeans in India. This was also reflected in the haplotype-based analysis, where one cluster of C2 segregated with these subpopulations (Fig S8). Even though the linkage disequilibrium-based method for admixture dating with ALDER revealed C2 and C3 to have had admixture history with all the contemporary west Eurasian groups from middle east, Europe and Caucasus (Fig S12A-F), the clue to a higher component of Middle Eastern ancestry was mirrored in the distal modelling where a high contribution from Bronze Age Namazga_CA like source group for both C2 and C3 (Fig 3A) was noted. This indicates that these two groups may have emerged from the same wave of migration with early Bronze Age Middle Eastern component. Even though the test for admixture failed in case of C3 with both modern and ancient reference data due to significant drift, a higher affinity with Namazga_CA is still evident, as only negative F3 statistics (supplementary table_2g) was observed with this group.

To understand the sources of ancestry, all the three Coorg groups were fitted in admixture graph topology for South Asian populations. This modelling suggested likely additional source of ancestry for C3 in both the best fitted graph topologies (Fig S3A-B). Accordingly, we propose that this group may actually harbour an ancient ancestry from a ghost population (unknown source as of date in our graph modelling approach), which alternatively may be a consequence of or account for the notable extent of drift. The population separation history using Relate also suggested that C3 separated much earlier than the typical timeframe for most of the Indo-European and Dravidian groups in India (Fig S13C). This possibility is amply supported by Y-STR network generated for both M780 and Z2125 markers, which were of completely independent Y chromosomal lineage. These may have arrived in India much earlier (based on their separation history) and with no admixture event with any R1a lineage of India thereafter (Fig 4A-B).

At this juncture, despite inhabiting distinct and isolated regions of South India, it may be relevant to discuss the shared genetic ancestry of C1 with Palliyar, a population group which has been reported earlier to have negligible Steppe contribution (Narasimhan, et al. 2019). This is discernible in their sharing of clade in TreeMix analysis with similar amount of drift (Fig 3) and also in fineSTRUCTURE tree (Fig S8-9). Of note, one cluster of C1 also showed affinity to Palliyar in the co-ancestry matrix as well (Fig S5). D-statistics to infer gene flow for C1 and Palliyar together also came up with negative statistics with West Eurasian and South Asian populations (supplementary table_1d) suggesting again that C1 and Palliyar belong to the same clade.

Population genomic studies with such a strong and unique population specific drift has not been reported till date in the Indian context. Previously, in a broader South Asian context, a single such instance related to Kalash population from Pakistan, which was considered as a genetic isolate because of the significant population specific drift has been reported (Ayub et al., 2015). Thus, the population-specific drift in C3, never before observed in Indian populations studied till date provides a good resource to study evolutionary phenomenon in natural populations using whole genome data in future. Furthermore, the presence of South Asian-specific haplogroup H in C3 with very high frequency (0.45) is quite surprising, owing to their Indo-European genetic composition as reflected in Admixture modelling approach (Fig2A-B). The only explanation may be that there was a much earlier migration and settlement of C3 in South Asia compared to other Indo-European migrations, assimilation of local group(s) and either lesser or complete absence of later admixture with any Indian population. Secondly, in an earlier study it was proposed that due to a significantly higher level of ASI ancestry in Palliyar like groups, they can be used as proxy for ASI in future studies (Narasimhan, et al. 2019). On the same note, C1 which has been witnessed to be a sister clade of Palliyar with significant similarities may also be used as a proxy for ASI in South Asian studies.

Finally, this study presents the first ever insightful evidence for an ancient origin and unique genetic architecture of the Coorg. The study findings unambiguously show that the present-day socio-culturally homogenous Coorg population is actually comprised of three distinct genetically heterogeneous clusters with all three population signatures dating back to early Bronze Age. One group (C1) was demonstrated to be a sister clade of the geographically distant Palliyar population with ancient ASI ancestry and had immense population-specific drift, indicative of their presence in India since antiquity. Another group (C3) also showed considerable degree of isolation which interfered with all the statistical analyses. However, it is evident that a) this group had a much higher contribution of ancient Bronze-Age Middle Eastern ancestry; b) they had diverged and separated much before mature-Bronze Age which stands in stark contrast to their Indian subcontinental neighbours; and c) an unknown ghost population might have at some point of time been the source of ancestry for this group but this remains unclear. Conversely, the C2 group was the most similar to its Indo-European contemporaries in terms of genetic landscape and population separation (mature Bronze-age). Yet, it stood distinct from its surrounding sub-populations because of its likely admixture with C3 and possessing a higher ancient Middle Eastern ancestry like that of the Nairs. This lends support to the early opinion of O’ Connor in his Memoir of Codagu 1870 where he mentions that the land was held by aborigines with feudal tenure (pt2, pg. 30); that he also referred to as a division of Malayalam Nairs (pg. 38-39). C1 is an old population and an integral part of the present-day Coorg population and yet shows a weak genetic affinity to C2 and C3. Taken together, it may be surmised that C1 (native) and C2 and C3 (neighbouring) sub-populations are all early Bronze Age and isolated independently at different points in time and space but may have eventually converged geographically and admixed genetically and/or socio-culturally and became the founding population of the current day Coorgs. A comparison of IBD scores relative to either Finnish or Ashkenazi Jews among the three Coorg groups clearly indicates that C1 with highest IBD score may be more susceptible to recessive genetic disorder. However, the negligible load of autosomal recessive disorders in this rather small (< 0.3 million) population practising family exogamy but caste endogamy may lend further support to such a contribution of diverse gene pools from the three groups through a later admixture among these geographically closely spaced populations.

Overall conclusion of the present study is that, the Coorg population has a complex genetic sub structuring with three distinct groups socio-culturally indistinguishable but having distinct genetic composition. We have also observed enormous genetic drift in Coorg3, which was not found among any Indian population on ANI-ASI cline till date, except Onge which is a population with history of migration much earlier than the major ANI-ASI admixture event. This group also requires an additional ghost population as an admixture edge in modelling approach, but may alternatively account for higher amount of genetic drift. An interesting history of bottleneck was also observed in Coorg1 along with comparatively higher distribution of RoH and higher IBD score. This group is quite similar in genetic composition and drift to a geographically distant group Palliyar and can alternatively be well suited as a proxy for Ancestral South Indian (ASI) population in modelling approaches.

## Material and Methods

### Study Subjects

Coorgs are a very small population group comprised of less than 0.3 million individuals, belonging to approximately 1200 extended families who follow endogamy. A representative sample set comprising of ∼10-12% of these families with only one male member from any generation per family was recruited for this study (n=144) with informed consent. and IEC. All individuals were healthy (self-reported) and in the age range of 25-70 years. About 5.0 ml of venous blood was drawn from each participant with their informed written consent. DNA was isolated using the standard phenol-chloroform method and used for subsequent genetic analysis.

### Genetic analysis

#### Genotyping

Genotyping of autosomal, mitochondrial, and Y chromosomal markers was performed as described below.

#### Autosomal markers

A subset of the samples (n=70) was genotyped using Affymetrix Axiom GW Human Origin Array for 633,994 SNPs as per the manufacturer’s specifications through a commercial facility (Imperial Life Sciences, Gurugram, Haryana).

#### Quality Control

The dataset was merged with published DNA dataset of contemporary Indian populations (Reich, et al. 2009; Moorjani, et al. 2013; Mallick, et al. 2016; Nakatsuka, et al. 2017) after filtering for missingness using Plink 1.9 (Chang, et al. 2015), and only autosomal markers on 22 chromosomes having genotyping call rate > 99% and minor allele frequency > 1% were included. Dataset was further pruned by removing individuals with first-degree and second-degree relatedness utilizing KING-robust (Manichaikul, et al. 2010) feature implemented in Plink2 (Chang, et al. 2015). After all filtering, final merged dataset comprised of 968 modern individuals genotyped at 380,944 SNPs.

In order to minimize the effect of background LD in PCA (Patterson, et al. 2006) and ADMIXTURE (Alexander, et al. 2009) like analysis, the markers were further thinned by removing SNPs in strong LD (r2 > 0.4, window of 200 SNPs, sliding window of 25 SNPs at a time) using Plink 1.9 (Chang, et al. 2015). For all the analysis with ancient DNA, Coorg samples were merged with west Eurasian autosomal DNA published datasets of 765 individuals with relevance to the incumbent sample set (Meyer, et al. 2012; Fu, et al. 2014; Pickrell, et al. 2014; Raghavan, et al. 2014; Allentoft, et al. 2015; Haak, et al. 2015; Broushaki, et al. 2016; Gallego-Llorente, et al. 2016; Lazaridis, et al. 2016; Yang, et al. 2017; Damgaard, et al. 2018; Narasimhan, et al. 2019). In this merged dataset, missingness criteria of geno > 0.7 was applied to include only those individuals covered at at least 70% of sites resulting into 968 individuals covered at 442230 sites.

#### mtDNA markers

Mitochondrial DNA of all samples were PCR amplified using a set of 24 sets of primer (Rieder, et al. 1998) followed by Sanger sequencing.

#### Y-chromosomal markers

Genotyping of all samples for a total of 18 Y-chromosomal binary markers to determine haplogroups was performed. PCR-amplified amplicons were sequenced using ABI 3730 automated Genetic Analyzer. Y-STR typing for 17 markers was done using ampFLSTR™ Yfiler™ PCR amplification kit.

## Data Analyses

### Autosomal

#### Principal Component Analysis

Principal Component analysis (PCA) was performed on the merged dataset of modern Eurasian using the *smartpca* package implemented in EIGENSOFT 7.2.1(Patterson, et al. 2006) with default settings. The first two components were plotted to infer genetic variability.

#### ADMIXTURE Analysis

Model-based clustering algorithm ADMIXTURE (Alexander, et al. 2009) was run to infer ancestral genomic components in Coorg population inferred from the PCA performed. Cross validation was run 25 times for 12 ancestral clusters (K=2 to K=12) (Fig S1). Lowest CV error parameter was obtained at K = 4 and was used for downstream analysis.

#### Maximum Likelihood tree construction

A maximum likelihood (ML) tree was constructed for the merged dataset comprising of modern South Asian populations and the Coorgs with TreeMix v.1.12 using LD blocks of 500 SNPs grouped together and Onge as an outgroup.

#### Runs of Homozygosity

Runs of Homozygosity (RoH) analysis was performed using PLINK v1.9 (Chang, et al. 2015) with three homozygous windows of 1000kb, 2500kb and 5000kb with minimum 50 consecutive SNPs.

#### IBD score calculation

IBD scores for the three Coorg groups relative to Finnish population was calculated with same pipeline as used in our earlier study (Nakatsuka, et al. 2017). SHAPEIT version 4.2.2 (Delaneau, et al. 2019) for phasing the genotype data and Refined-IBD tool (Browning and Browning 2013) for IBD detection was used. Then slightly modified R script from Nathan et al. 2017 was used for IBD score calculation.

#### F3-statistics and D-statistics

*qp3Pop* and *qpDstat* implementation of ADMIXTOOLS (Patterson, et al. 2012) package was utilised to calculate Admixture F3 statistics and D-statistics, respectively. To infer gene flow from modern Eurasians in three groups of Coorg populations F3 statistics was used in the form of F3 (X, Palliyar; Coorg1/Coorg2/Coorg3), where X is any modern west Eurasian or south Asian population and Palliyar was used as a proxy for ASI ancestry. D-statistics was calculated in the form of F4 (X, Coorg1/Coorg2/Coorg3, French; Yoruba) and F4 (X, Coorg1/Coorg2/Coorg3, Palliyar; Yoruba) to infer the extent of West Eurasian specific and Ancestral south Asian specific gene flow into three groups of Coorg population.

### ALDER

In order to gain insight into time scale of admixture events, if any, in the Coorg population, ALDER (Loh, et al. 2013) was run, which uses exponential decay of linkage disequilibrium to infer the approximate time of admixture. For this, different contemporary West Eurasian populations and Juang (source of South Asian component) were used as putative admixing source populations.

#### CHROMOPAINTER and FineStructure

Haplotype-based approach implemented in CHROMOPAINTER (Lawson, et al. 2012) and FineStructure (Lawson, et al. 2012) was used to derive co-ancestry matrix and fine scale population clustering, respectively. Data was first phased with SHAPEIT4 using default parameters, followed by CHROMOPAINTER run to infer co-ancestry matrix, first by performing 10 Expectation-Maximization (EM) iteration with 5 randomly selected chromosomes with a subset of individuals to infer global mutation rate (µ) and switch rate parameters (Ne). Then the main algorithm was run with 22 chromosomes with all the individuals to derive the co-ancestry matrix. This matrix was used by FineStructure to derive clustering using a probability model by applying Markov chain Monte Carlo (MCMC) procedure and then inferring hierarchical tree by merging all clusters with least change in posterior probability. For the run 500,000 burn-in iterations and 1,000,000 subsequent iterations were used, and the results stored from every 10,000th iteration.

#### Proximal and Distal modelling with ancient DNA

*qpAdm* in the ADMIXTOOLS 2 (Maier, et al. 2022) package in R was used to estimate proportions of ancient ancestral components in a test population (Coorg1/Coorg2/Coorg3) derived from a set of N source population groups having shared drift with a set of reference populations. Distal and Proximal modelling of admixture was performed using early-Bronze Age aDNA source groups and Bronze Age proximal sources, respectively. In distal modelling AHG, Namazga_CA and Steppe_MLBA were used as source groups, with Ethiopia_4500BP_published.SG, ANE, Natufian, PPNB, Dai.DG, EEHG, Ganj_Dareh_N and WEHG as references. In proximal modelling AHG, Indus_Periphery and Steppe_MLBA were used as source groups and Ethiopia_4500BP_published.SG, Ganj_Dareh_N, EEHG, PPNB, Dai.DG, Anatolia_N, WEHG were taken as references. Fitted admixture graph topology were obtained with *qpGraph* function of ADMIXTOOLS 2 (Maier, et al. 2022) using automated graph exploration with *find_graph* for three Coorg groups using modern and ancient Eurasians as reference. *qpGraph* was further used to model Coorg2 and Coorg3 along with Kalash and Gujjar as a mixture of ANI and ASI ancestry, using the model (YRI, (Coorg2/Coorg3/Kalash/Gujjar, (Georgians, ANI)), [(ASI, Onge])) proposed by Moorjani et al (Moorjani, et al. 2013). This method was used earlier in Nakatsuka et al. (2017) (Nakatsuka, et al. 2017) to estimate the strength of founder effect in Indian populations by measuring post-admixture drift.

##### mtDNA

Sequences were assembled with the reference sequence rCRS (Andrews, et al. 1999) using AutoAssembler. Variations observed were used to assign the haplogroup using phylotree build 17 (van Oven and Kayser 2009) and Haplogrep2 (Weissensteiner, et al. 2016).

##### Y-chromosomal

Sequences were compared with reference to mark the variations and assign the haplogroups. In order to estimate population divergence, Weir-Cockerham’s F_st_ was measured using R package SambaR (de Jong, et al. 2021) utilising the same pruned dataset as used in PCA and Admixture. For the Y STR data analysis, the Y-LineageTracker (Chen, et al. 2021) tool was used. Population separation history was inferred using genome-wide genealogy based approach implemented in Relate package (Speidel, et al. 2019) using haplotype data phased in SHAPEIT4.2 (Delaneau, et al. 2019).

## Supporting information

Supplementary Sheet_1

Supplimentary Sheet_2

## Acknowledgements

We thank all the study participants who volunteered in this study. B.K.T designed the study and collected the samples on-site. Kiran Sran carried out initial haplogrouping work of uniparental markers. A.M performed further haplogrouping work. A.M and L.K performed analysis of autosomal data. A.M and L.K and B.K.T wrote the first draft of the paper. All authors contributed to and have approved the final manuscript. The authors have declared that there no conflicts of interest in relation to the subject of this study. Grateful thanks to Ms Deepika Singh, UDSC for DNA isolations and Mr. Jagmohan Chattai, CCMB for his assistance with PCR-sequencing for Y-STR markers. We deeply acknowledge the critical inputs on the manuscript provided by Mr. Mookonda Nitin Kushalappa, independent researcher and author. Council of Scientific and Industrial Research (CSIR) Fellowship to L.K, J.C Bose fellowship (#SR/S2/JCB44/2011 & 2016) to B.K.T and JC Bose Fellowship to K.T, from Science and Engineering Research Board, New Delhi; One time grant from university grants Commission, New Delhi to BKT are gratefully acknowledged. Computational facilities provided by Central Instrumentation Facility, University of Delhi South Campus; Infrastructure support by the Department of Science and Technology, New Delhi, for FIST and DU-DST PURSE programmes to the Department of Genetics, UDSC, are gratefully acknowledged.

## Notes

### Competing Interest Statement

The authors have declared no competing interest.

