## Supplementary_Figures and Text for "Genetic affinities and sub-structuring in Coorg population of Southern India"

**Supplementary Figure 1.** Mean Admixture cross validation error plotted at different K values from K=2 to K=12.

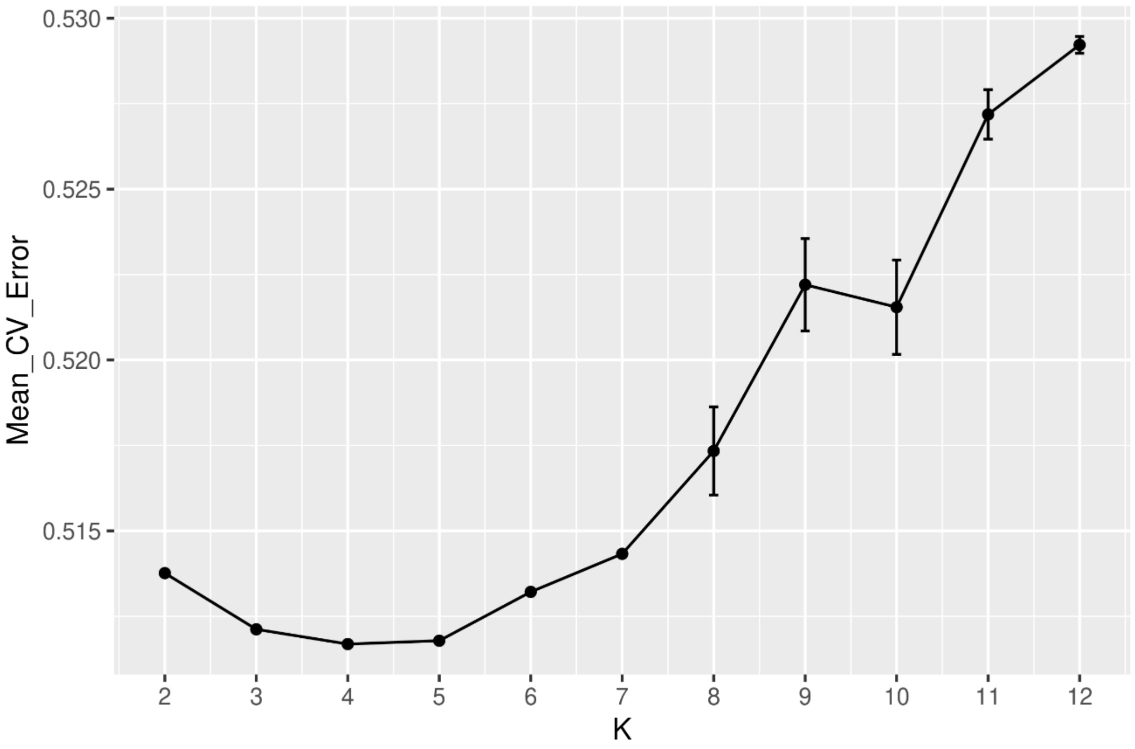

**Supplementary Figure 2.** Scatterplot of comparison of affinity of Coorg groups with modern Eurasian population for west Eurasian ancestry in the form F4(X, Coorg1/Coorg2/Coorg3; French, Yoruba) against ASI ancestry in the form F4(X, Coorg1/Coorg2/Coorg3; Palliyar, Yoruba), where X is any other west Eurasian or south Asian population.

(A) Scatterplot for Coorg1 (B) Scatterplot for Coorg2 (C) Scatterplot for Coorg3

**A.**

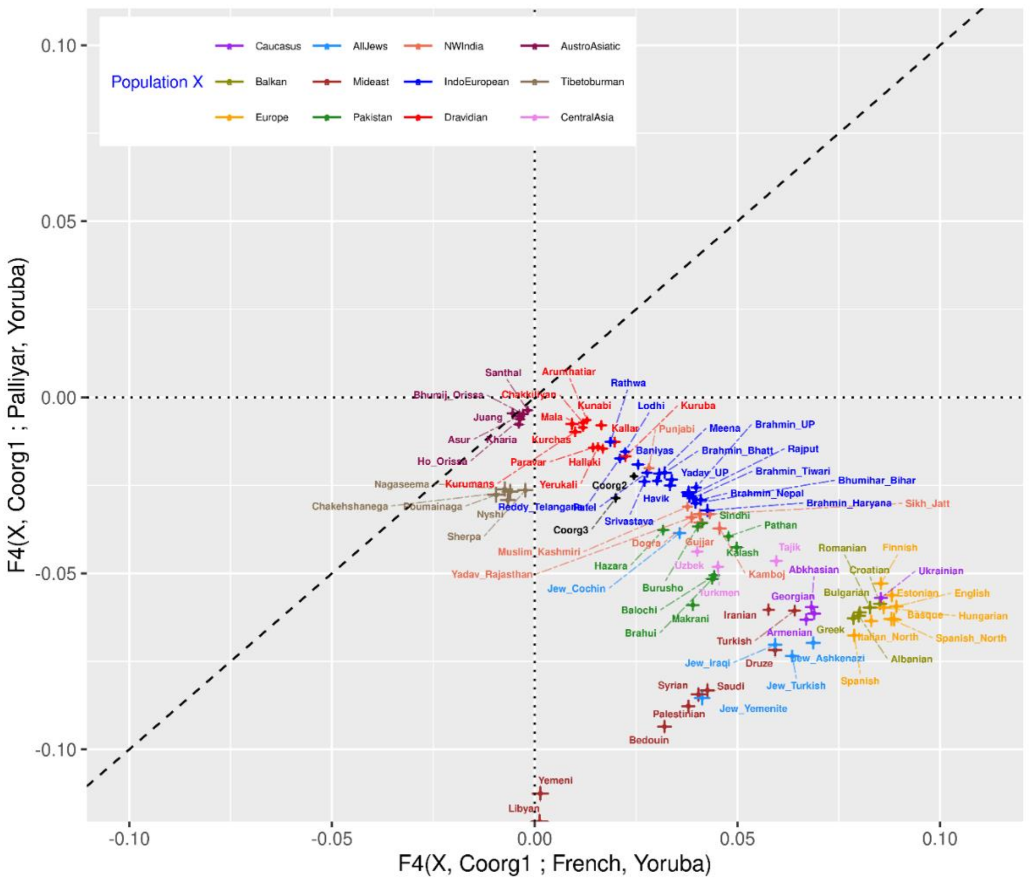

**B.**

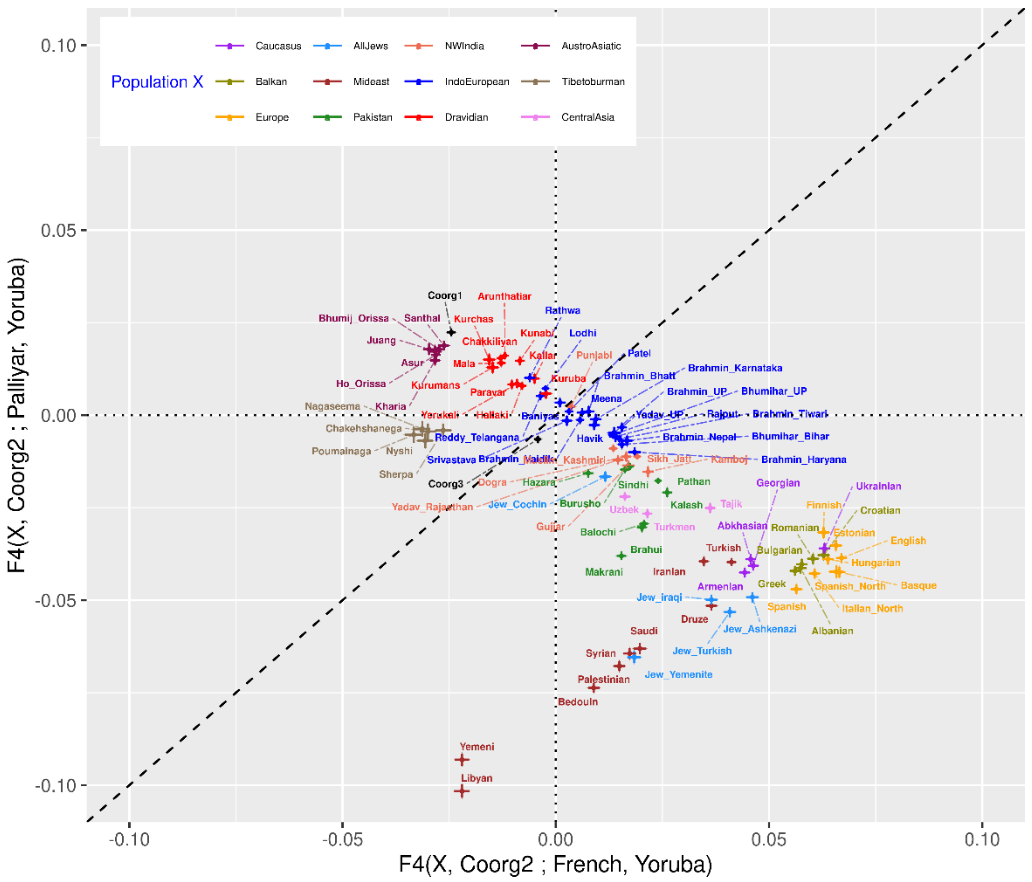

Population X
Caucasus
AllJews
NWIndia
AustroAsiatic
Balkan
Mideast
IndoEuropean
Tibetoburman
Europe
Pakistan
Dravidian
CentralAsia
0.10
0.05
0.00
-0.05
-0.10
-0.10
-0.05
0.00
0.05
0.10
F4(X, Coorg2 ; Palliyar, Yoruba)
F4(X, Coorg2 ; French, Yoruba)


**C.**

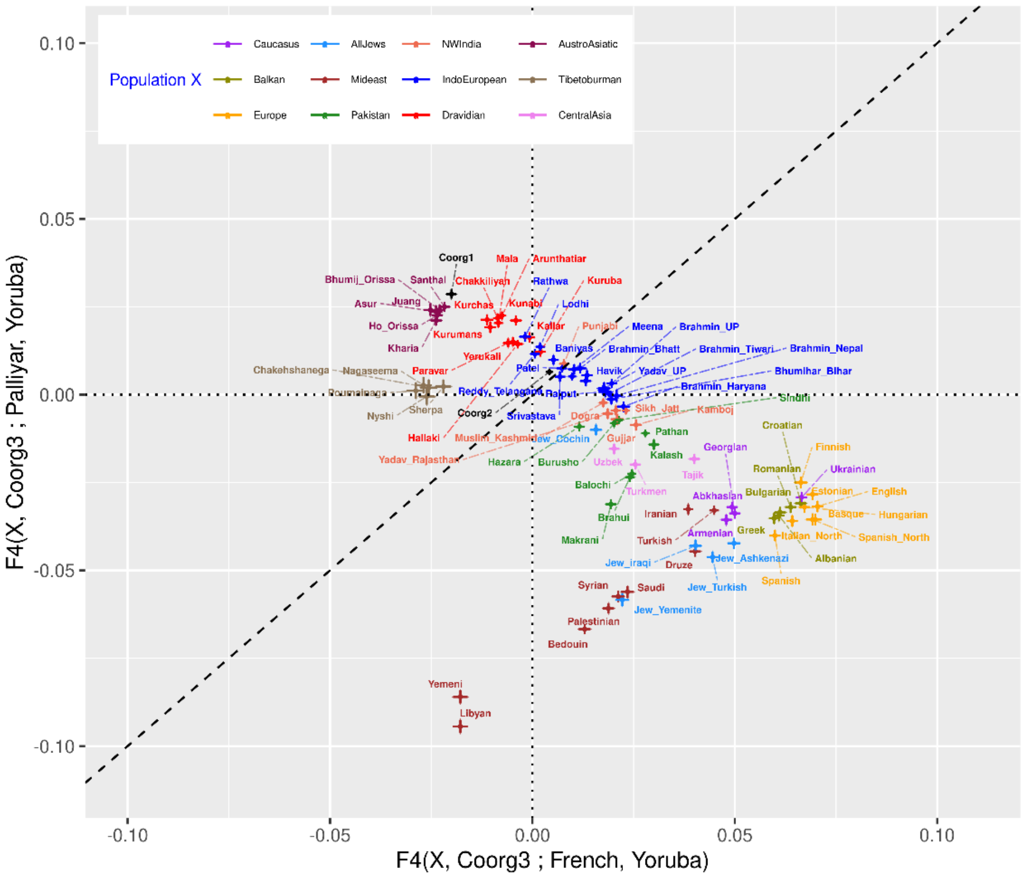

Population X
Caucasus
AllJews
NWIndia
AustroAsiatic
Balkan
Mideast
IndoEuropean
Tibetoburman
Europe
Pakistan
Dravidian
CentralAsia
0.10
0.05
0.00
-0.05
-0.10
-0.10
-0.05
0.00
0.05
0.10
F4(X, Coorg3 ; Palliyar, Yoruba)
F4(X, Coorg3 ; French, Yoruba)

**Supplementary Figure 3.** Admixture graph model for three Coorg groups using ancient source of ancestry.

(A) Fitted graph topology (Graph1) for Coorg1, Coorg2 and Coorg3 with likelihood score 2.032829, showing that Coorg3 additional admixture edge with 1% ancestry from unknown source.

(B) Fitted graph topology (Graph2) for Coorg1, Coorg2 and Coorg3 with likelihood score 2.417081, showing that Coorg3 additional admixture edge with 9% ancestry from unknown source.

**A.**

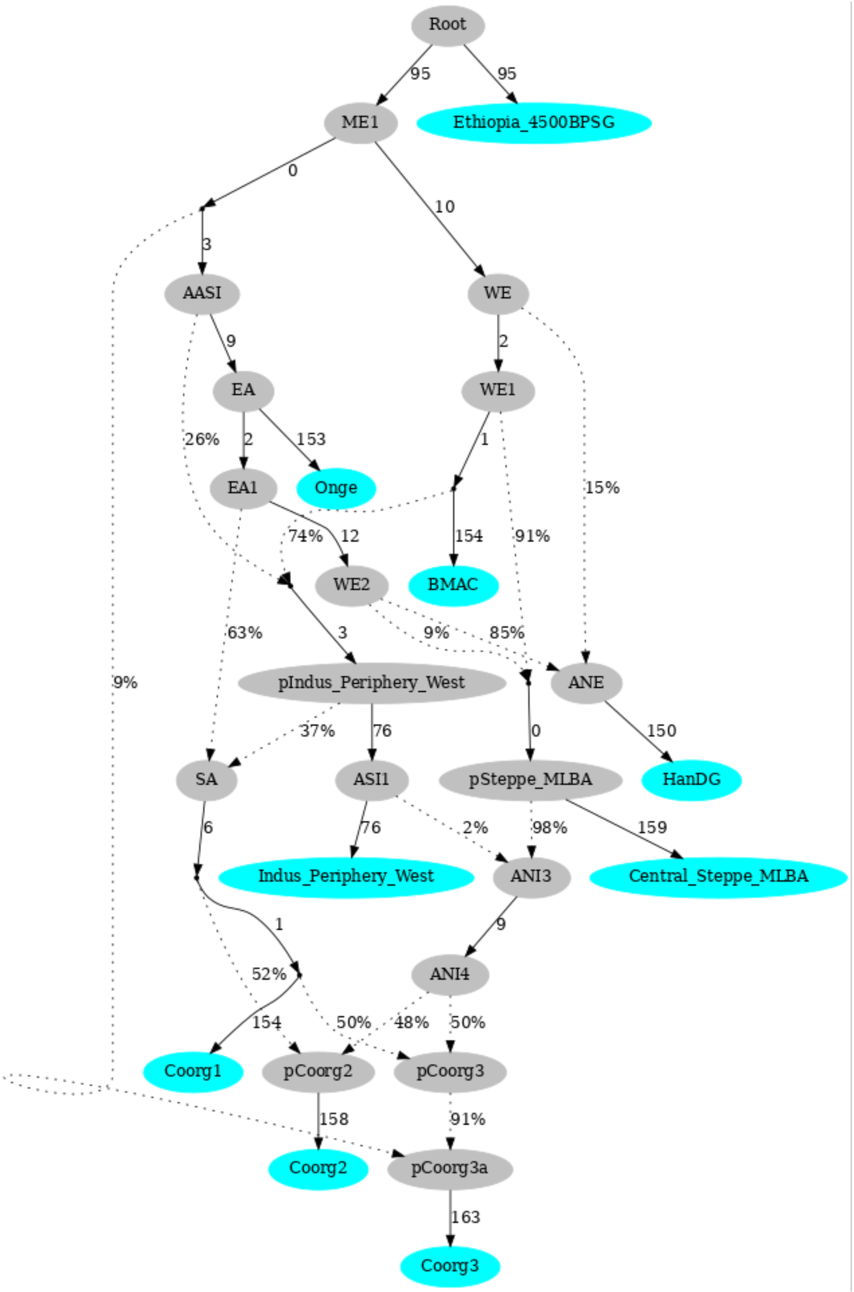

**B.**

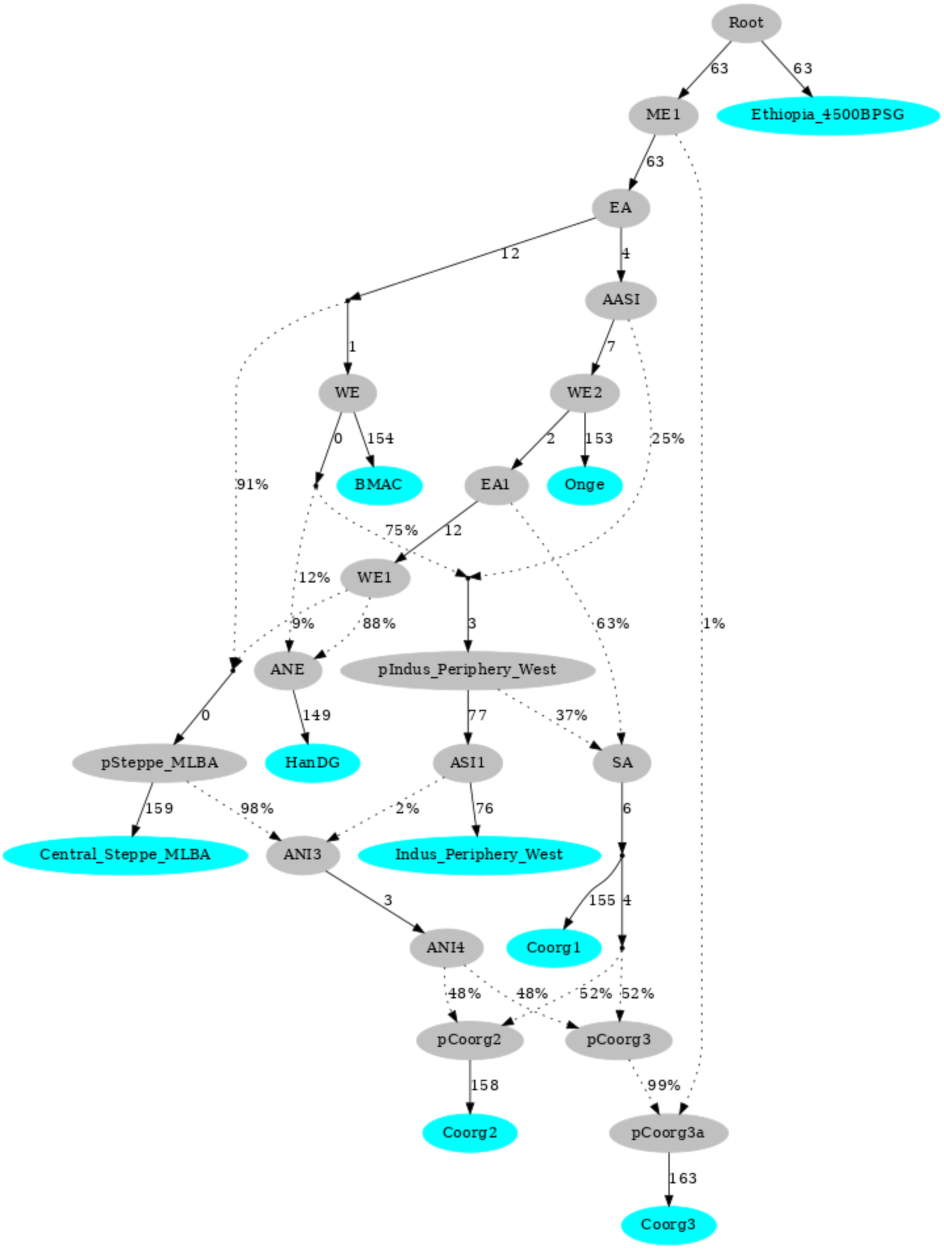

Root
63
63
ME1
Ethiopia_4500BPSG
63
EA
12
4
AASI
1
7
WE
WE2
0
154
2
153
25%
91%
BMAC
EA1
Onge
75%
12
12%
WE1
9%
88%
3
63%
1%
ANE
pIndus_Periphery_West
0
149
77
37%
pSteppe_MLBA
HanDG
ASI1
SA
159
98%
2%
76
6
Central_Steppe_MLBA
ANI3
Indus_Periphery_West
3
155
4
ANI4
Coorg1
48%
48%
52%
52%
pCoorg2
pCoorg3
158
99%
Coorg2
pCoorg3a
163
Coorg3

**Supplementary Figure 4. Genome wide distribution of Wier-Cockerham Fst value for autosomal SNPs.** (A) Coorg1 with Dravidian, (B) Coorg1 with European, (C) Coorg1 with Middle East, (D) Coorg2 with Dravidian, (E) Coorg2 with European, (F) Coorg2 with Middle East, (G) Coorg3 with Dravidian, (H) Coorg3 with European, (I) Coorg3 with Middle East.

**A.**

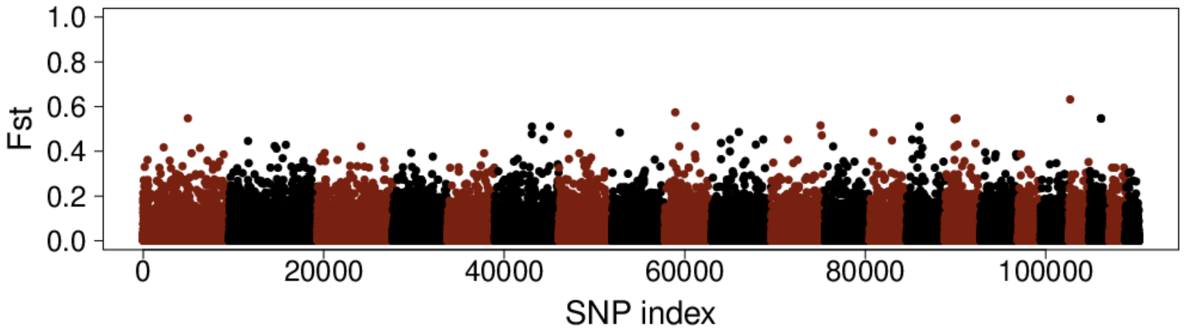


**B.**

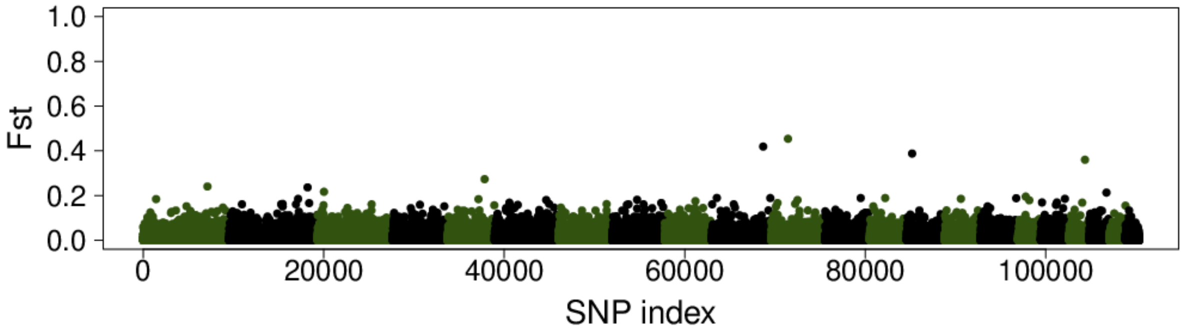


**C.**

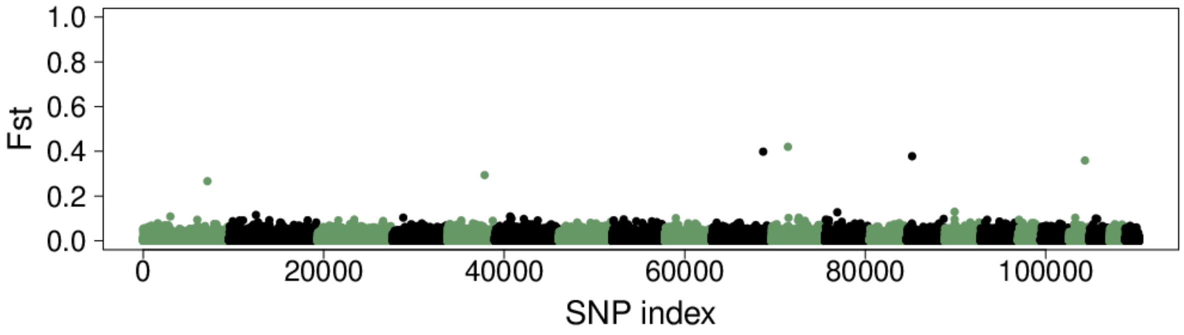

D.

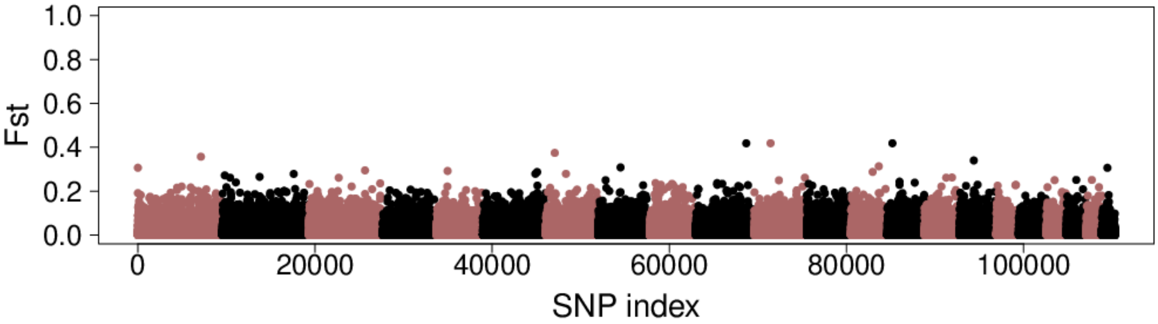
Fst
1.0
0.8
0.6
0.4
0.2
0.0
0
20000
40000
60000
80000
100000
SNP index

E.

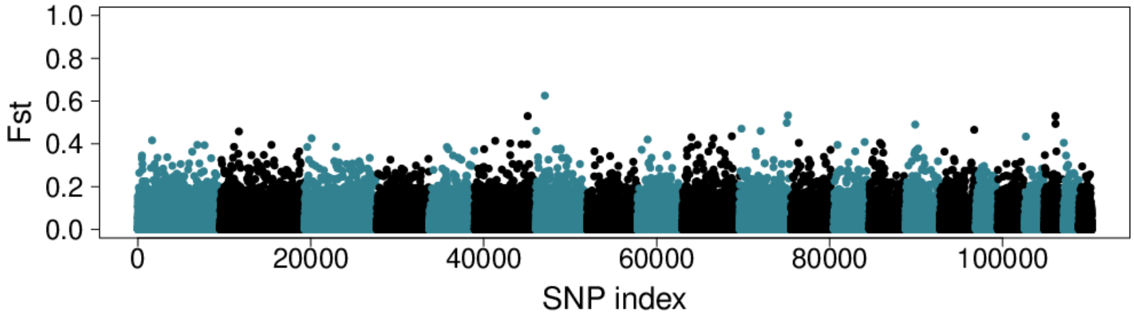
Fst
1.0
0.8
0.6
0.4
0.2
0.0
0
20000
40000
60000
80000
100000
SNP index

F.

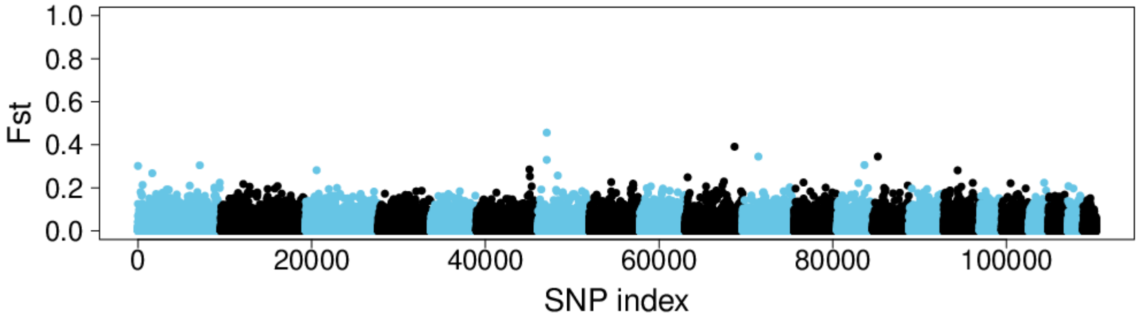
Fst
1.0
0.8
0.6
0.4
0.2
0.0
0
20000
40000
60000
80000
100000
SNP index

G.

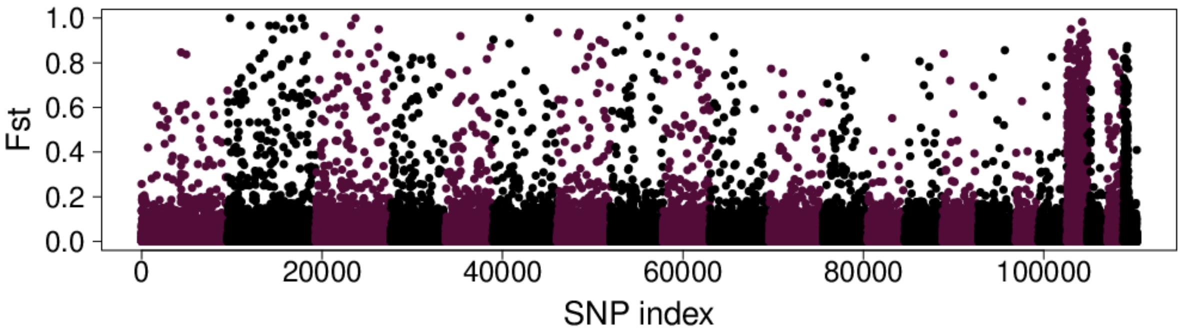

Fst
1.0
0.8
0.6
0.4
0.2
0.0
0
20000
40000
60000
80000
100000
SNP index


H.

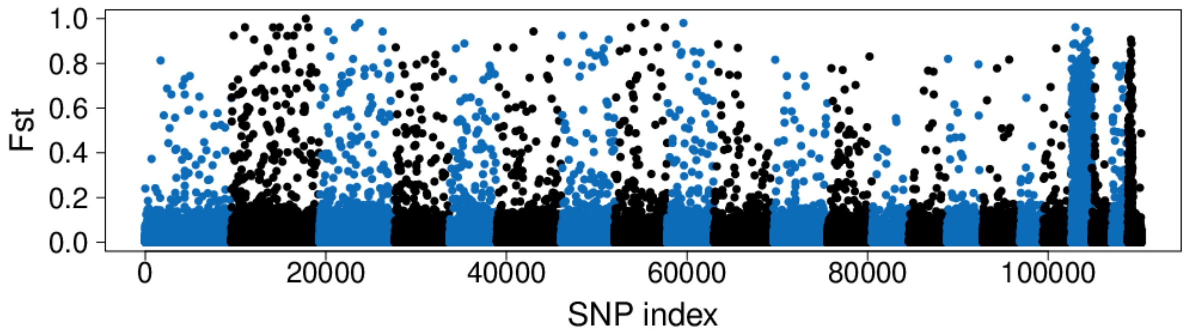

Fst
1.0
0.8
0.6
0.4
0.2
0.0
0
20000
40000
60000
80000
100000
SNP index


I.

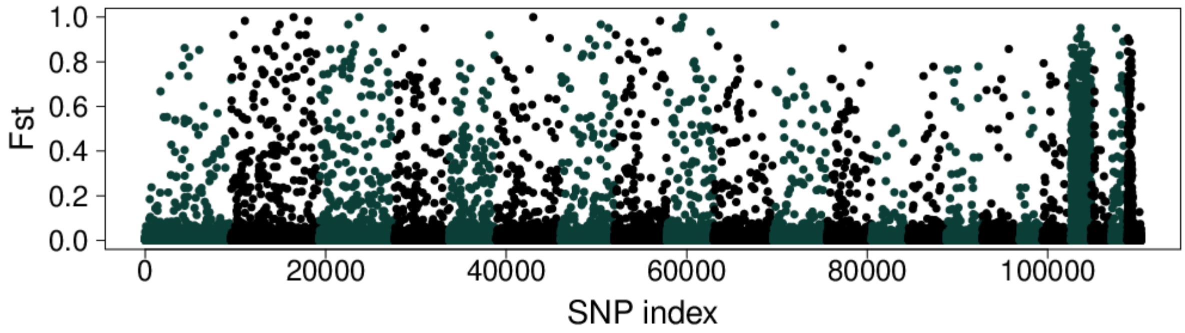

Fst
1.0
0.8
0.6
0.4
0.2
0.0
0
20000
40000
60000
80000
100000
SNP index

**Supplementary Figure 5.** Inter-population pairwise Wier-Cockerham Fst distance matrix for three groups of Coorg with Europe, Middle East, Indo-Europeans and Dravidian from India.

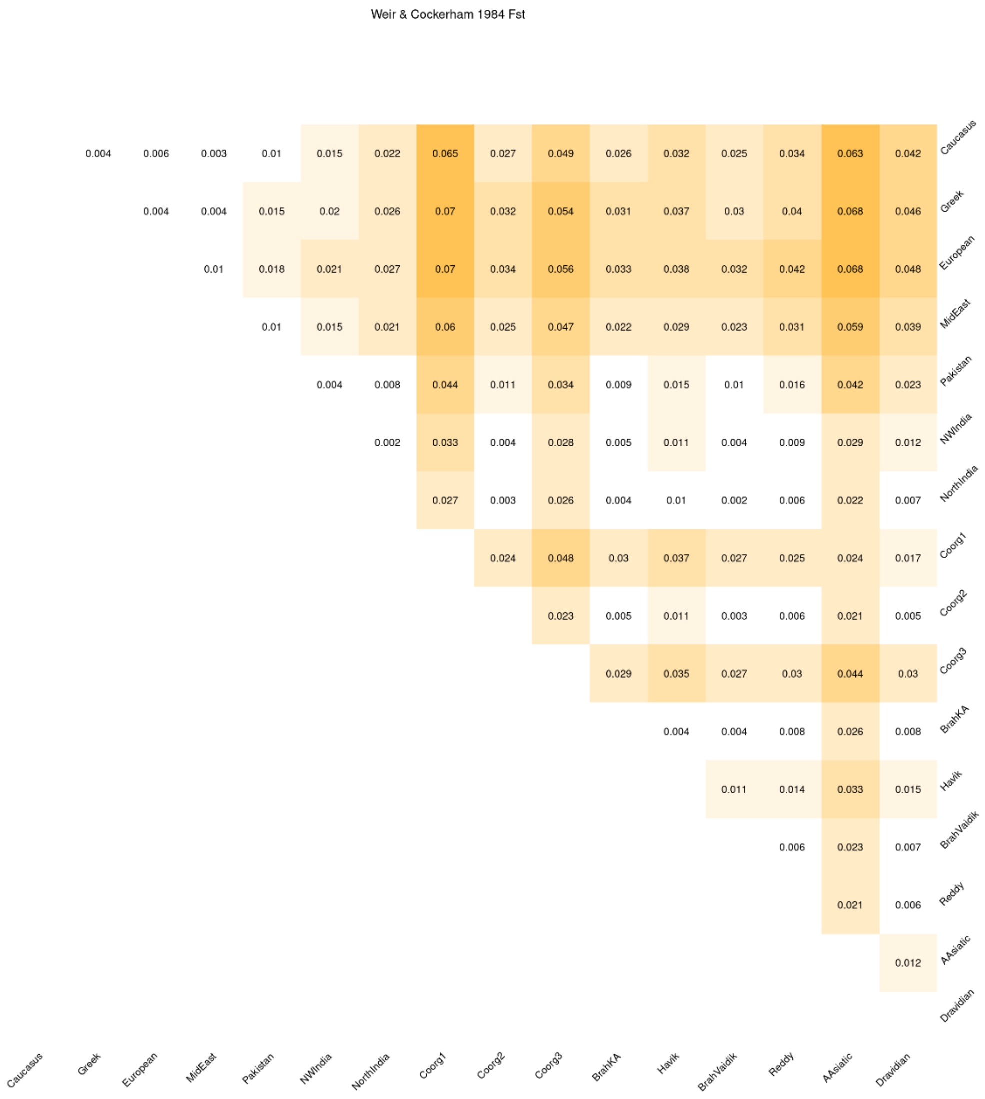

**Supplementary Figure 6.** qpGraph method for the estimation of population specific drift among four groups of **(A)**Coorg2, **(B)**Coorg3, **(C)**Kalash and **(D)**Gujjar. R=root; OoA=Out of Africa; ASA=Ancestral South Asian; ASI=Ancestral Southern Indian; AWE=Ancestral West Eurasian; ANI=Ancestral North Indian; APOP=Ancestral Indian group. Branch lengths in the units of FST x 1,000.

A.

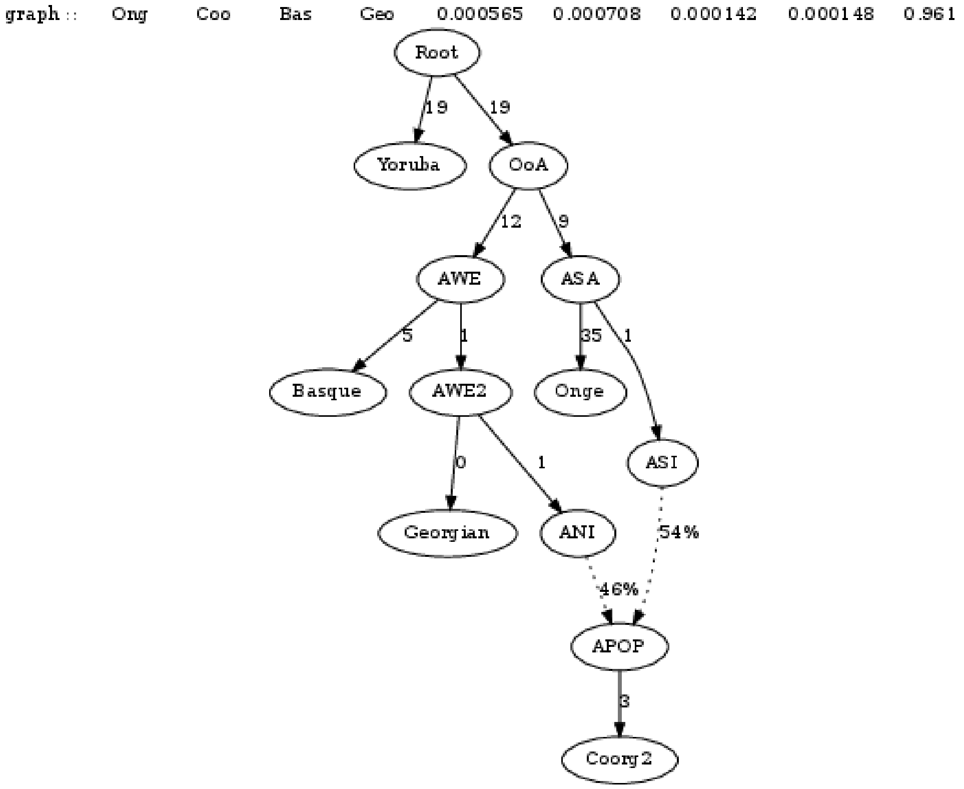

**B.**

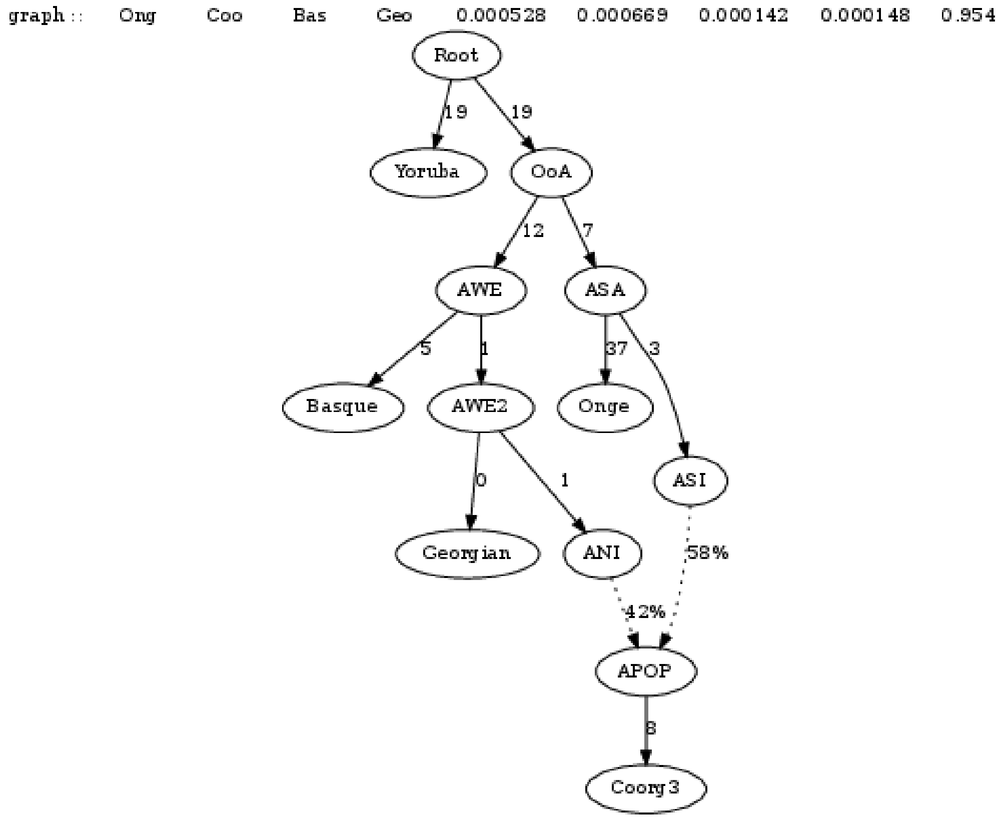
graph :: Ong Coo Bas Geo 0.000528 0.000669 0.000142 0.000148 0.954
Root
19
19
Yoruba
OoA
12
7
AWE
ASA
5
1
37
3
Basque
AWE2
Onge
0
1
ASI
Georgian
ANI
58%
42%
APOP
8
Coorg3

C.

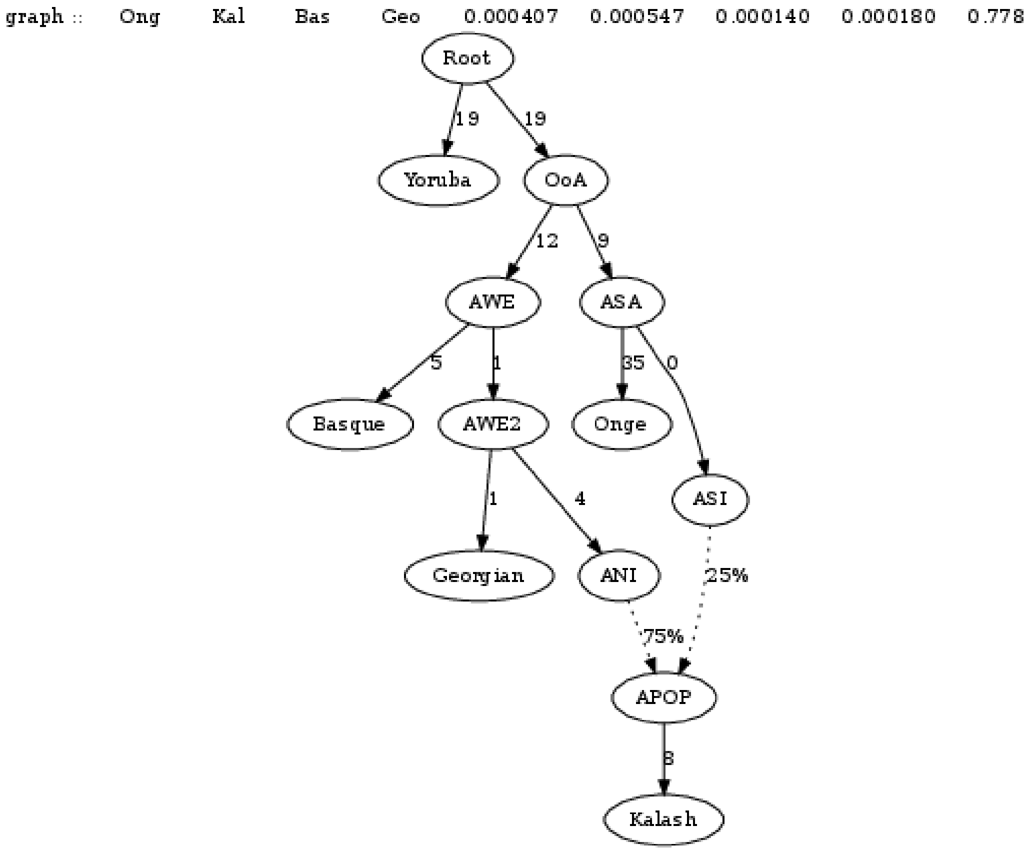
graph :: Ong Kal Bas Geo 0.000407 0.000547 0.000140 0.000180 0.778
Root
19
19
Yoruba
OoA
12
9
AWE
ASA
5
1
35
0
Basque
AWE2
Onge
ASI
1
4
Georgian
ANI
25%
75%
APOP
8
Kalash

**D.**

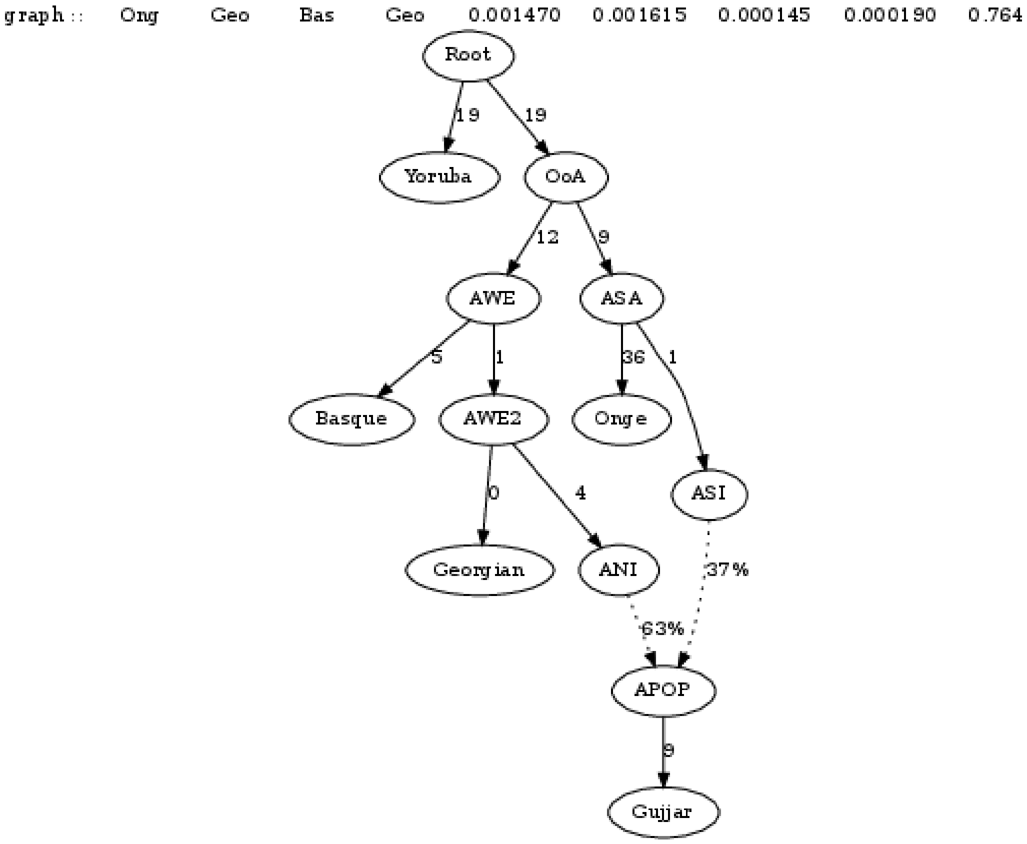
graph :: Ong Geo Bas Geo 0.001470 0.001615 0.000145 0.000190 0.764
Root
19
19
Yoruba
OoA
12
9
AWE
ASA
5
1
36
1
Basque
AWE2
Onge
0
4
ASI
Georgian
ANI
37%
63%
APOP
9
Gujjar

**Supplementary Figure 7.** Biplot of Principal component analysis using coancestry matrix generated by haplotype-based method implemented chromopainter.

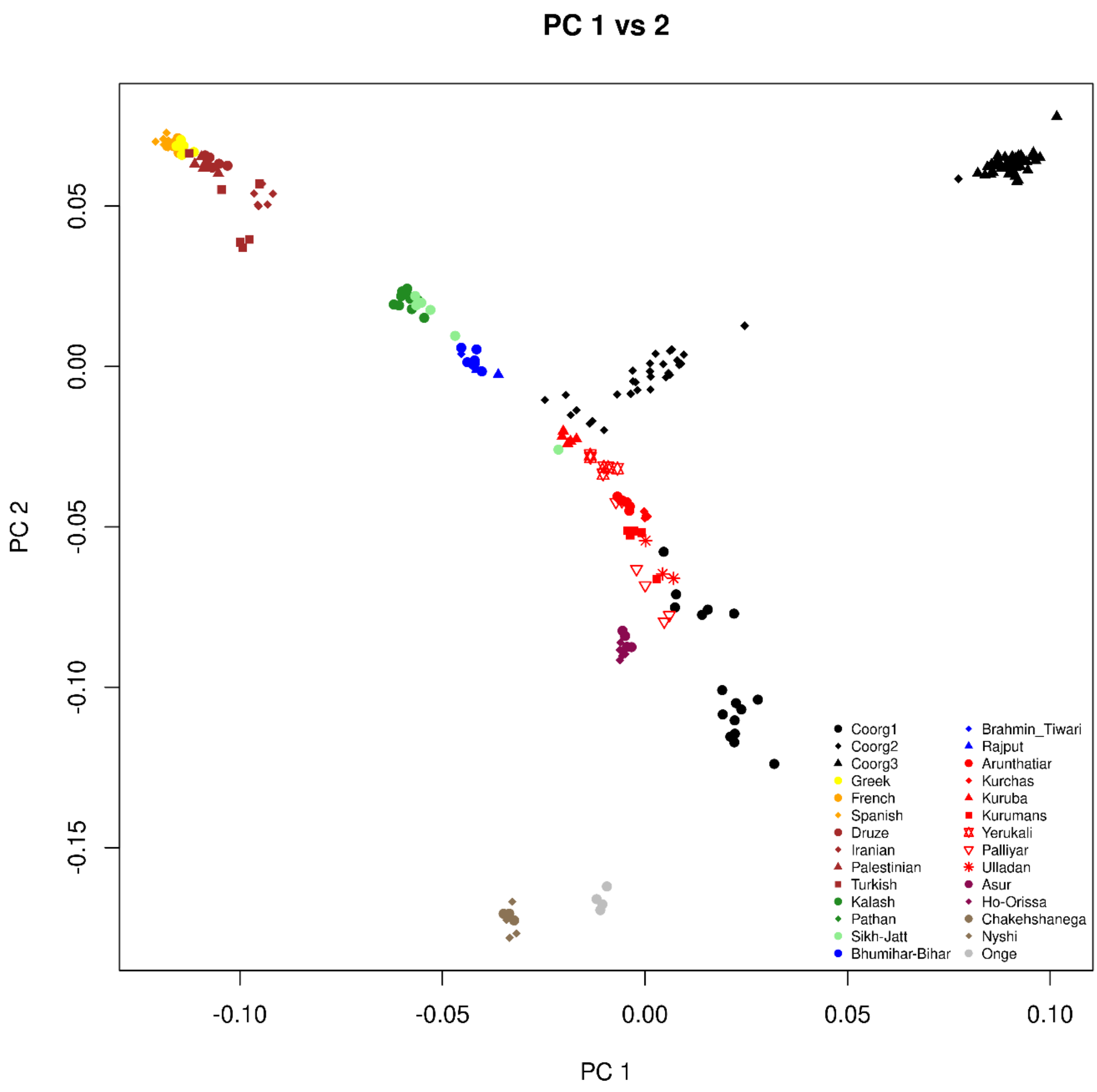

**Supplementary Figure 8.** Placement of three groups of Coorg individuals among 44 worldwide clades in population dendrogram generated by fineStructure. Individuals of group 3 making separate clade (details of individuals in supplementary fig 9).

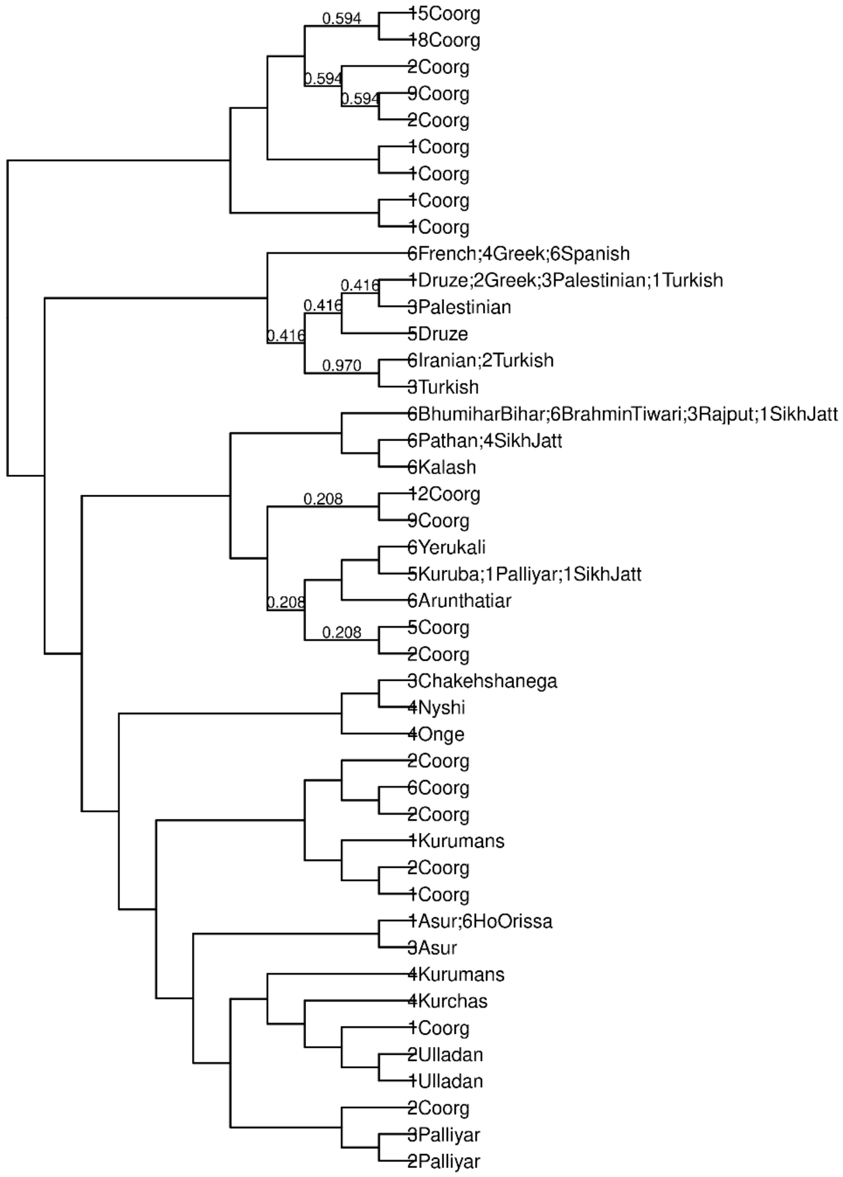

**Supplementary Figure 9.** Full population dendrogram generated by fineStructure with individual annotations. Coorg group 3 is comprised of individuals from Coorg46 to Coorg94, all are present in a separate clade from all other Eurasians. Two individuals of group 2 are also included in this separate clade

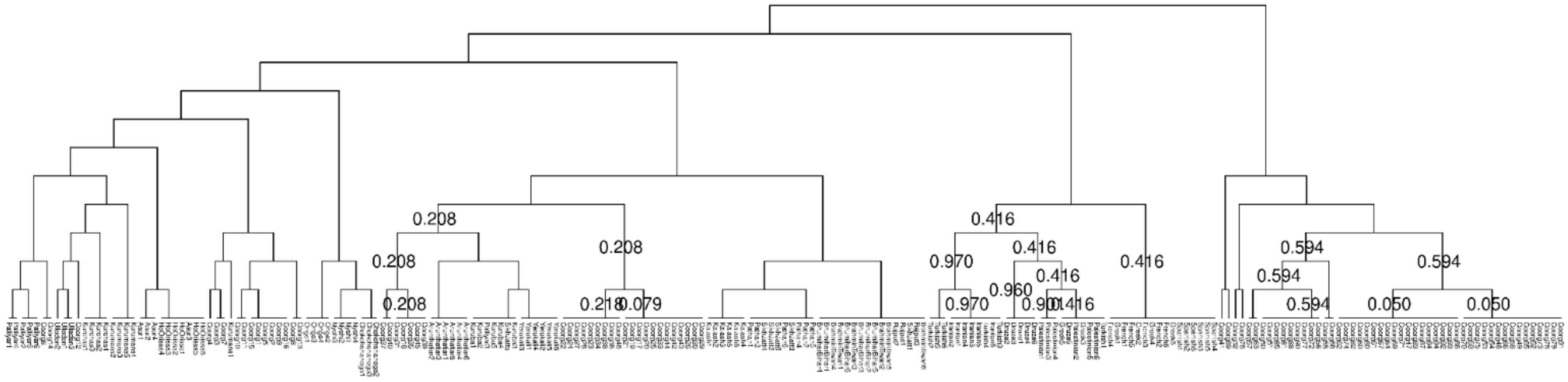

**Supplementary Figure 10.** Scatter plot of the average lengths of Runs of Homozygosity (RoH) against average number of RoH segments for three Coorg groups and other South Asians using three different windows of (A) 1000 kb, (B) 2500 kb and (C) 5000 kb.

**A.**

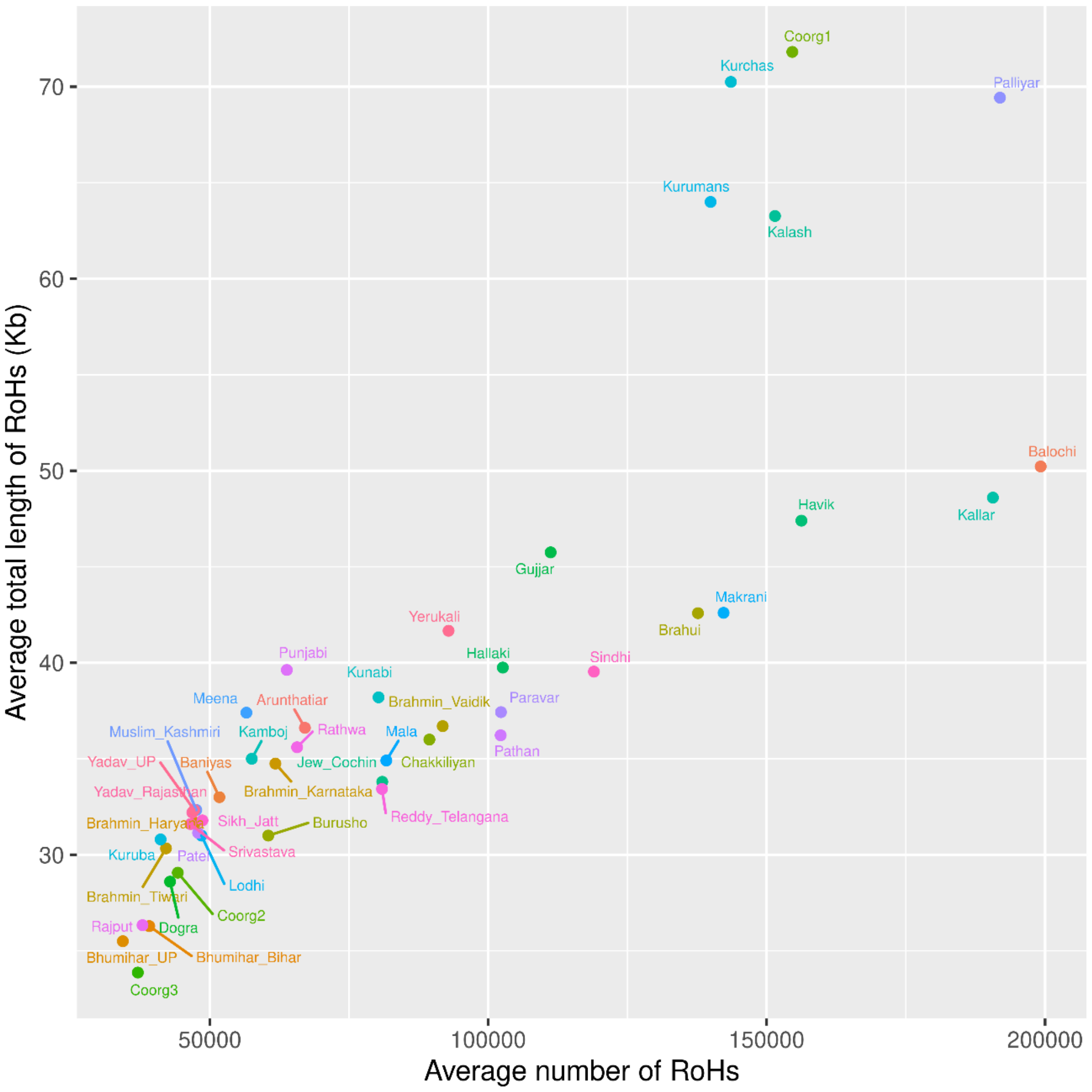

**B.**

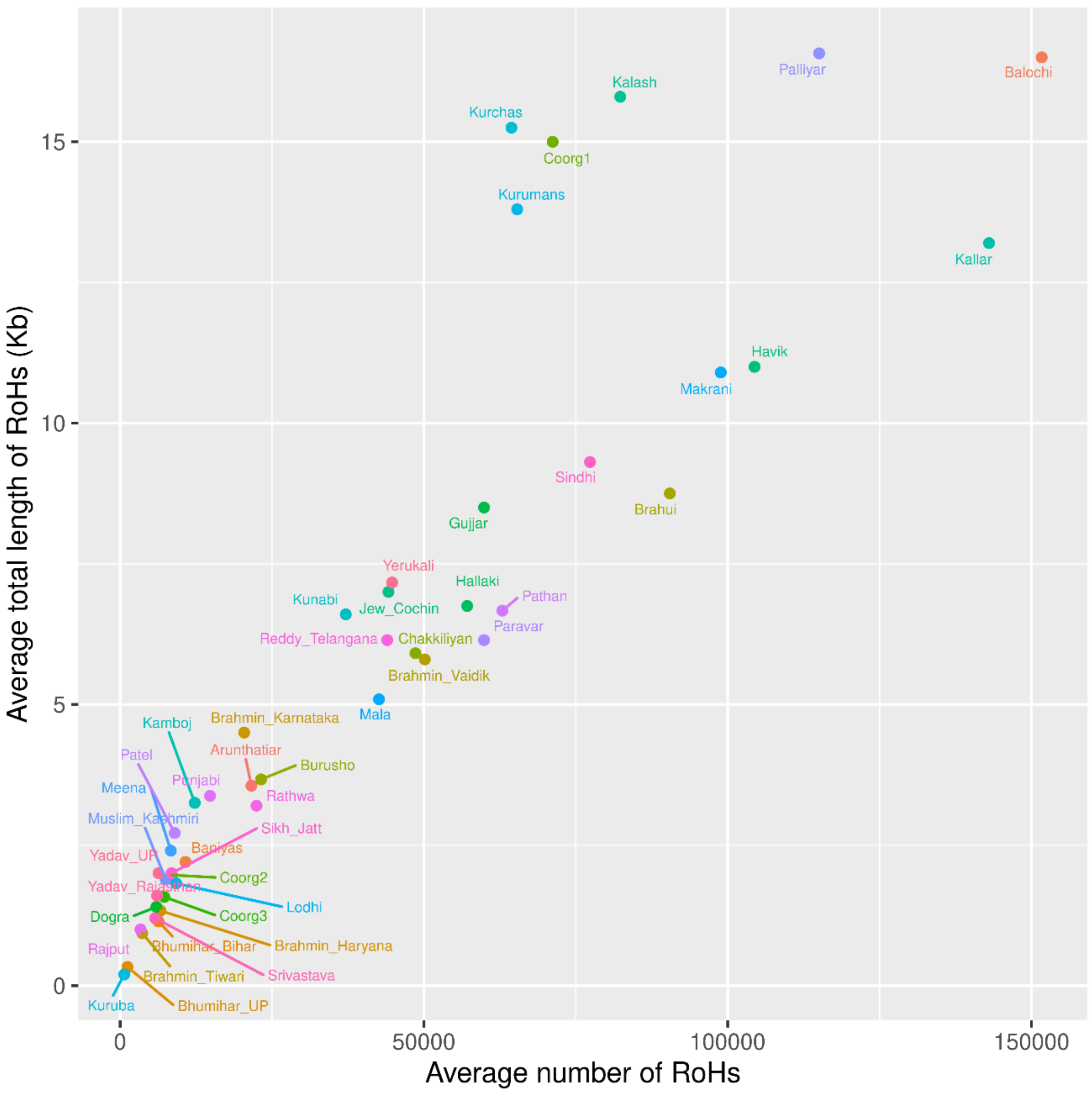

Average total length of RoHs (Kb)
Average number of RoHs
0
5
10
15
0
50000
100000
150000
Palliyar
Balochi
Kalash
Kurchas
Coorg1
Kurumans
Kallar
Havik
Makrani
Sindhi
Brahui
Gujjar
Yerukali
Hallaki
Pathan
Kunabi
Jew_Cochin
Paravar
Reddy_Telangana
Chakkiliyan
Brahmin_Vaidik
Mala
Kamboj
Brahmin_Karnataka
Patel
Arunthatiar
Burusho
Meena
Punjabi
Rathwa
Muslim_Kashmiri
Sikh_Jatt
Yadav_UP
Baniyas
Yadav_Rajasthan
Coorg2
Lodhi
Dogra
Coorg3
Rajput
Bhumihar_Bihar
Brahmin_Haryana
Brahmin_Tiwari
Srivastava
Kuruba
Bhumihar_UP

**C.**

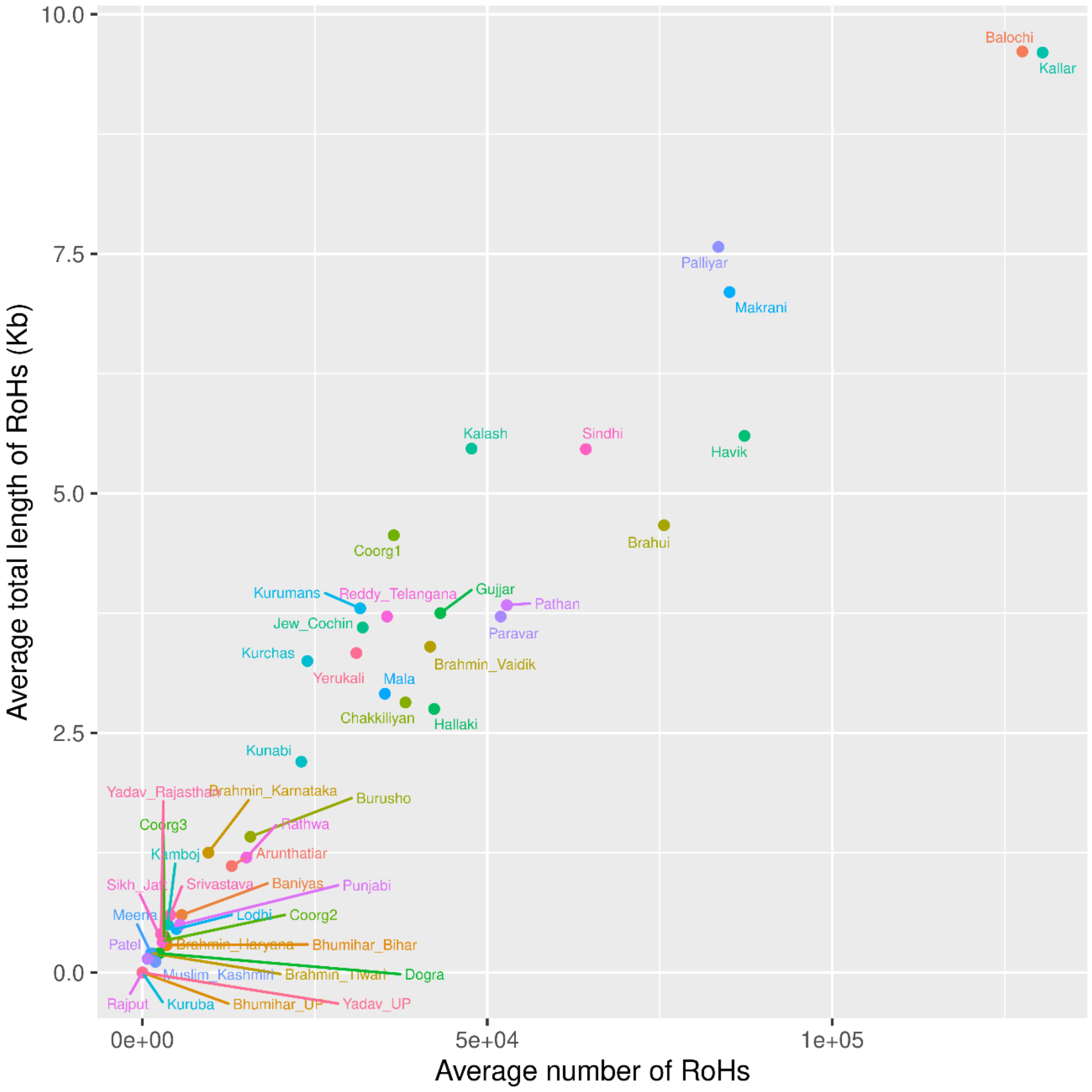
Average total length of RoHs (Kb)
Average number of RoHs
10.0
7.5
5.0
2.5
0.0
0e+00
5e+04
1e+05
Balochi
Kallar
Palliyar
Makrani
Kalash
Sindhi
Havik
Brahui
Coorg1
Kurumans
Reddy_Telangana
Gujjar
Pathan
Jew_Cochin
Paravar
Kurchas
Brahmin_Vaidik
Yerukali
Mala
Chakkiliyan
Hallaki
Kunabi
Yadav_Rajasthan
Brahmin_Karnataka
Burusho
Coorg3
Rathwa
Kamboj
Arunthatiar
Sikh_Jat
Srivastava
Baniyas
Punjabi
Meena
Lodhi
Coorg2
Patel
Brahmin_Haryana
Bhumihar_Bihar
Muslim_Kashmiri
Brahmin_Tiwari
Dogra
Rajput
Kuruba
Bhumihar_UP
Yadav_UP

**Supplementary Figure 11.** IBD score of three Coorg groups along with populations from India with significantely higher IBD score in relative to Finnish population.

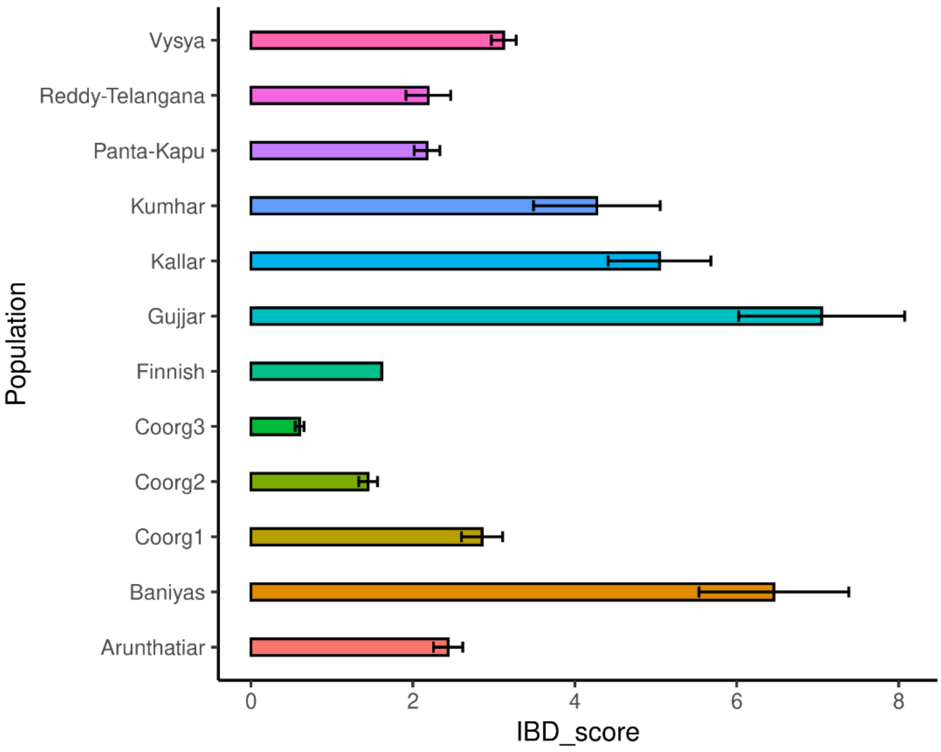

**Supplementary Figure 12.** Weighted LD decay curve Coorg2 and Coorg3 groups with different Eurasian source groups.

(A) LD decay curve of Coorg2 with Druze vs Juang, (B) LD decay curve of Coorg2 with French vs Juang weights, (C) LD decay curve of Coorg2 with Georgian vs Juang weights, (D) LD decay curve of Coorg3 with Druze vs Juang weights, (E) LD decay curve of Coorg3 with French vs Juang weights, (F) LD decay curve of Coorg3 with Georgian vs Juang weights.

**A.**

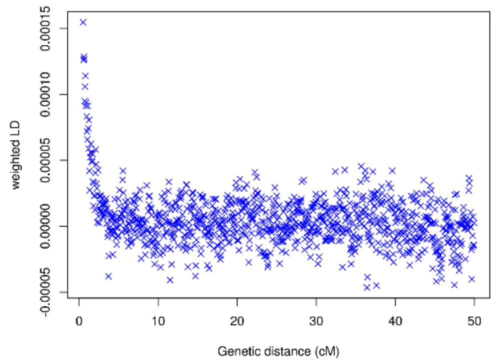


**B.**

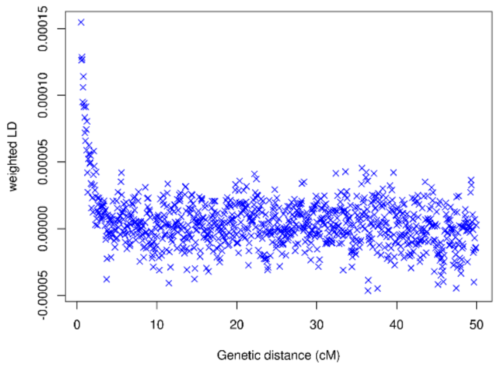


**C.**

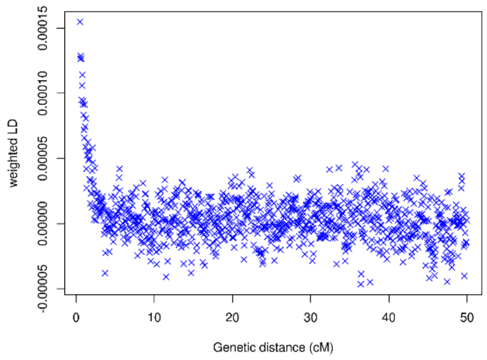


**D.**

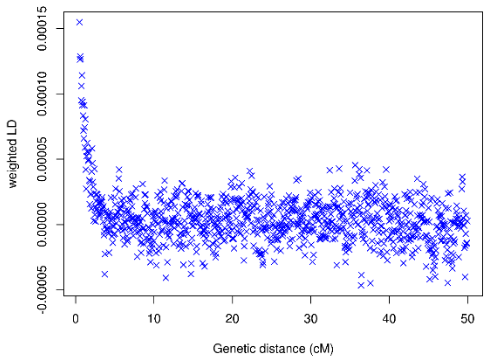


**E.**

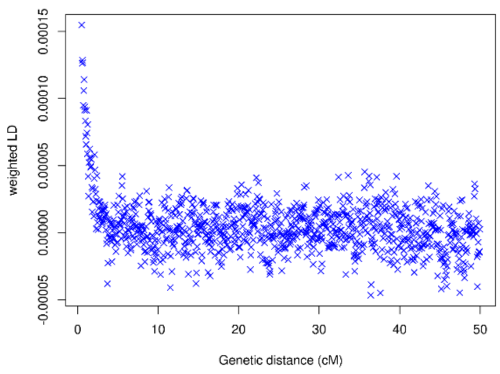


**F.**

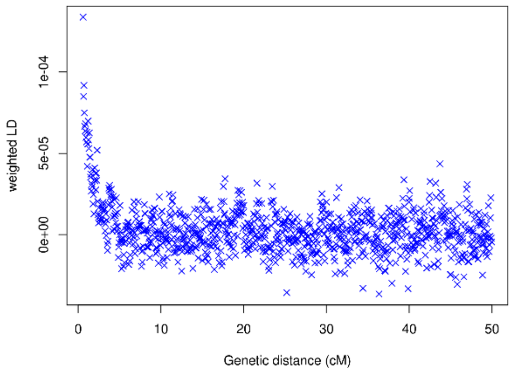

**Supplementary Figure 13.** Effective population size and population separation history

(A) population separation history for Coorg1, (B) population separation history for Coorg2, (C) population separation history for Coorg3.

**A.**

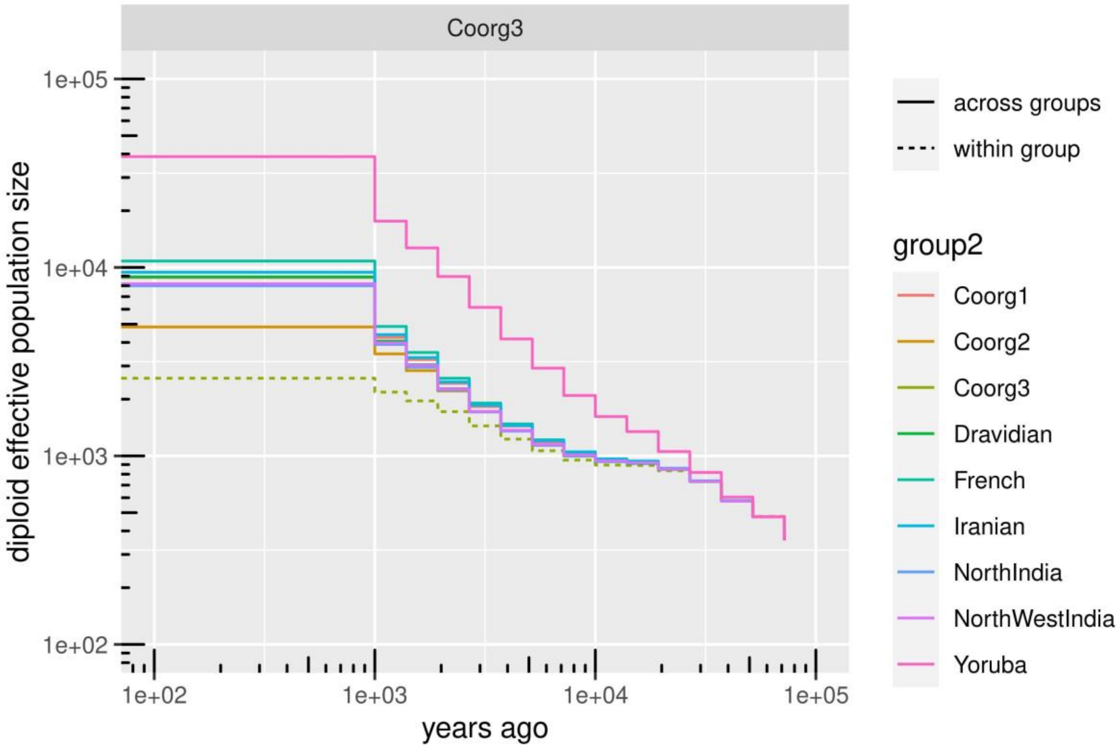

**B.**

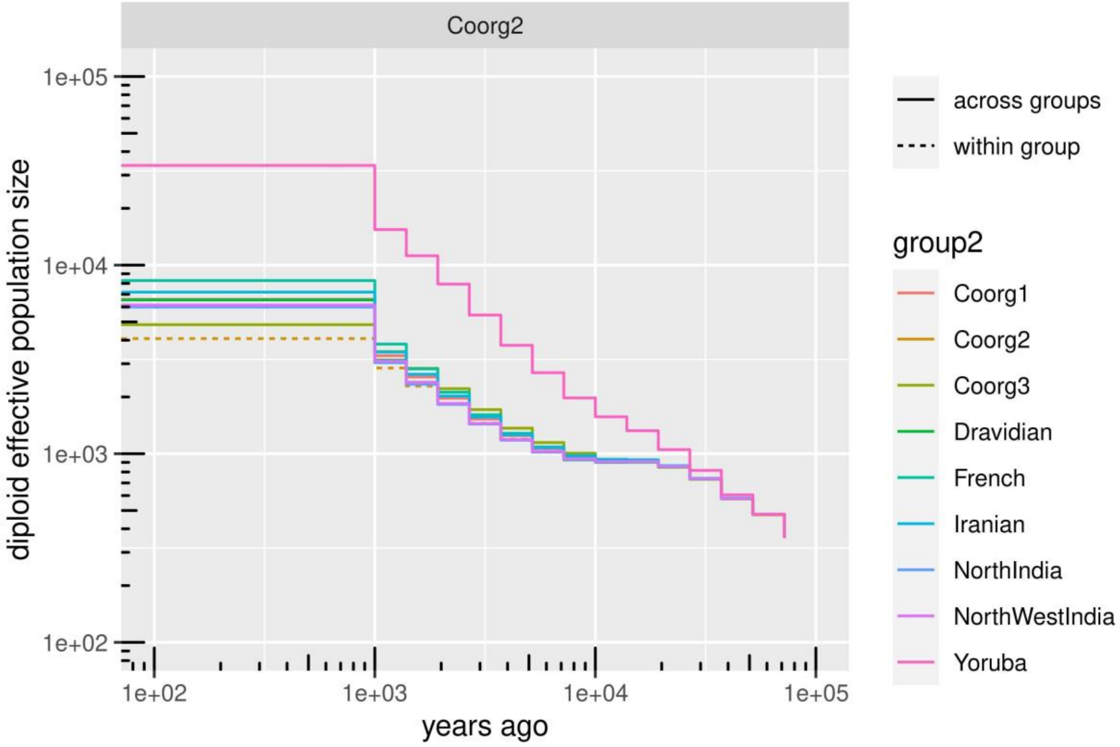
Coorg2
1e+05
1e+04
1e+03
1e+02
diploid effective population size
1e+02
1e+03
1e+04
1e+05
years ago
across groups
within group
group2
Coorg1
Coorg2
Coorg3
Dravidian
French
Iranian
NorthIndia
NorthWestIndia
Yoruba

**C.**

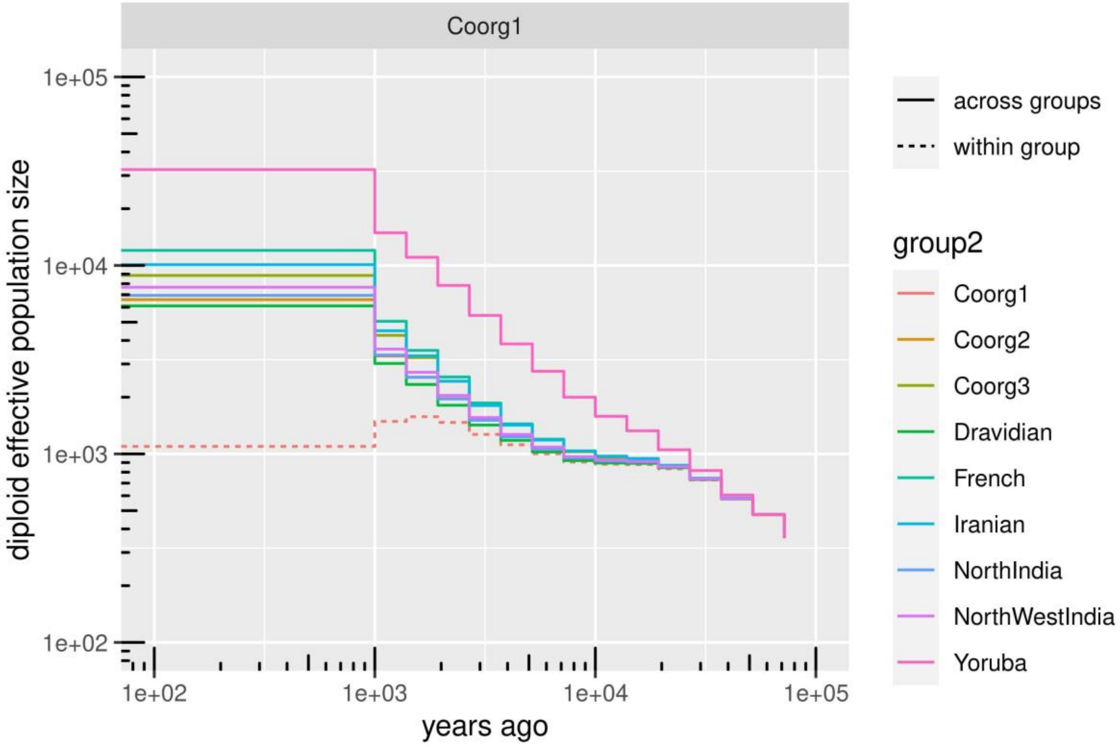
Coorg1
1e+05
1e+04
1e+03
1e+02
diploid effective population size
1e+02
1e+03
1e+04
1e+05
years ago
across groups
within group
group2
Coorg1
Coorg2
Coorg3
Dravidian
French
Iranian
NorthIndia
NorthWestIndia
Yoruba

### Supplementary Figure Legends

**Supplementary Figure 1.** Mean Admixture cross validation error plotted at different K values from K=3 to K=14.

**Supplementary Figure 2.** Scatterplot of comparison of affinity of Coorg groups with modern Eurasian population for west Eurasian ancestry in the form F4(X, Coorg1/Coorg2/Coorg3; French, Yoruba) against ASI ancestry in the form F4(X, Coorg1/Coorg2/Coorg3; Palliyar, Yoruba), where X is any other west Eurasian or south Asian population.

**Supplementary Figure 3.** Admixture graph model for three Coorg groups using ancient source of ancestry.

(A) Fitted graph topology (Graph1) for Coorg1, Coorg2 and Coorg3 with likelihood score 2.032829, showing that Coorg3 additional admixture edge with 1% ancestry from unknown source.

(B) Fitted graph topology (Graph2) for Coorg1, Coorg2 and Coorg3 with likelihood score 2.417081, showing that Coorg3 additional admixture edge with 9% ancestry from unknown source.

**Supplementary Figure 4. Genome wide distribution of Wier-Cockerham Fst value for autosomal SNPs.** (A) Coorg1 with Dravidian, (B) Coorg1 with European, (C) Coorg1 with Middle East, (D) Coorg2 with Dravidian, (E) Coorg2 with European, (F) Coorg2 with Middle East, (G) Coorg3 with Dravidian, (H) Coorg3 with European, (I) Coorg3 with Middle East.

**Supplementary Figure 5.** Inter-population pairwise Wier-Cockerham Fst distance matrix for three groups of Coorg with Europe, Middle East, Indo-Europeans and Dravidian from India.

**Supplementary Figure 6.** qpGraph method for the estimation of population specific drift among four groups of **(A)**Coorg2, **(B)**Coorg3, **(C)**Kalash and **(D)**Gujjar. R=root; OoA=Out of Africa; ASA=Ancestral South Asian; ASI=Ancestral Southern Indian; AWE=Ancestral West Eurasian; ANI=Ancestral North Indian; APOP=Ancestral Indian group. Branch lengths in the units of FST x 1,000.

Supplementary Figure *7*. Biplot of Principal component analysis using coancestry matrix generated by haplotype-based method implemented chromopainter.

**Supplementary Figure 8.** Placement of three groups of Coorg individuals among 44 worldwide clades in population dendrogram generated by fineStructure. Individuals of group 3 making separate clade (details of individuals in supplementary fig 9).

**Supplementary Figure 9.** Full population dendrogram generated by fineStructure with individual annotations. Coorg group 3 is comprised of individuals from Coorg46 to Coorg94, all are present in a separate clade from all other Eurasians. Two individuals of group 2 are also included in this separate clade

**Supplementary Figure 10.** Scatter plot of the average lengths of Runs of Homozygosity (RoH) against average number of RoH segments for three Coorg groups and other South Asians using three different windows of (A) 1000 kb, (B) 2500 kb and (C) 5000 kb.

**Supplementary Figure 11.** IBD score of three Coorg groups along with populations from India with significantely higher IBD score in relative to Finnish population.

**Supplementary Figure 12.** Weighted LD decay curve Coorg2 and Coorg3 groups with different Eurasian source groups.

(A) LD decay curve of Coorg2 with Druze vs Juang, (B) LD decay curve of Coorg2 with French vs Juang weights, (C) LD decay curve of Coorg2 with Georgian vs Juang weights, (D) LD decay curve of Coorg3 with Druze vs Juang weights, (E) LD decay curve of Coorg3 with French vs Juang weights, (F) LD decay curve of Coorg3 with Georgian vs Juang weights.

**Supplementary Figure 13.** Effective population size and population separation history

(A) population separation history for Coorg1, (B) population separation history for Coorg2, (C) population separation history for Coorg3.

### Supplementary Table Legends

#### Supplementary Sheet 1

**Supplementary Table 1a**. Admixture F3 statistics of Coorg3 as target, Palliyar (as proxy for ASI) as one source and various south Asian and west Eurasian populations as other source groups.

**Supplementary Table 1b.** Admixture F3 statistics of Coorg1 as target, Palliyar (as proxy for ASI) as one source and various south Asian and west Eurasian populations as other source groups.

**Supplementary Table 1c.** Admixture F3 statistics of Coorg group2 as target, Palliyar (as proxy for ASI) as one source and various south Asian and west Eurasian populations as other source groups.

**Supplementary Table 1d.** Dstatistics in the form F4(X, Coorg1; Palliyar, Yoruba), to show that Coorg2 and Palliyar belong to same clade, where population X is any south Asian or west Eurasian population.

**Supplementary Table 1e.** Dstatistics in the form F4(X, Coorg2; French, Yoruba), to infer west Eurasian affinity of Coorg2, where population X is any south Asian or west Eurasian population.

**Supplementary Table 1f.** Dstatistics in the form F4(X, Coorg3; French, Yoruba), to infer west Eurasian affinity of Coorg3, where population X is any south Asian or west Eurasian population.

**Supplementary Table 1g.** Output of Distal and Proximal admixture modelling with qpAdm for three Coorg groups and other Indo-European and Dravidian populations.

**Supplementary Table 1h.** Out of sample and bootstrap resample score for comparison of admixture graph1 and graph2.

**Supplementary Table 1i.** Results of IBD score calculation of three Coorg groups along with other Indian groups relative to Finnish population.

**Supplementary Table 1.** Details of ALDER run results with Coorg groups as targets, Juang as south Asian reference and different west Eurasian populations as other source groups.

(j) ALDER test outputs of Coorg3 as target population.

(k) ALDER test outputs of Coorg2 as target population.

(l) ALDER test outputs of Coorg1 as target population.

### Supplementary Sheet 2

**Supplementary Table 2a. Details of mtDNA and Y-DNA markers of Coorg1**

**Supplementary Table 2b. Details of mtDNA and Y-DNA markers of Coorg2**

**Supplementary Table 2c. Details of mtDNA and Y-DNA markers of Coorg3**

**Supplementary Table 2d. Details of mtDNA and Y-DNA markers for the complete dataset**
